# Pharmacological Inhibition of SLC33A1 Promotes Endoplasmic Reticulum Hyperoxidation and Induces Adaptive IRE1/XBP1s Signaling

**DOI:** 10.64898/2026.02.17.706344

**Authors:** Sergei Kutseikin, Maria Rafiq, Prerona Bora, Shanshan Liu, Rick A. Homan, Jeffrey T. Mindrebo, Matthew Holcomb, H. Michael Petrassi, Huang Qiu, Anastasiya Redkina, Justyna Sosna, Thanh-Trang Lee, Xiao Hu, Stefano Forli, Christopher G. Parker, Gabriel C. Lander, Kivanc Birsoy, Enrique Saez, R. Luke Wiseman

## Abstract

The endoplasmic reticulum (ER) transporter solute carrier family 33 member 1 (SLC33A1) has emerged as an attractive therapeutic target in etiologically diverse diseases, ranging from lung cancer to neurodegenerative disorders. Yet, no pharmacologic SLC33A1 modulators have been described. Here, we show that the small molecule IXA4, a highly selective activator of the adaptive IRE1/XBP1s signaling arm of the unfolded protein response (UPR), binds to SLC33A1 and inhibits its activity. Genetic depletion of *SLC33A1* phenocopies the selective induction of IRE1/XBP1s signaling brought about by IXA4 treatment. Chemoproteomic analyses and cryo-electron microscopy show that IXA4 binds SLC33A1 within the central channel to inhibit transport of its substrate metabolite(s). Binding of IXA4 to SLC33A1 leads to the accumulation of oxidized glutathione within the ER, hyperoxidizing the ER lumen and inducing activation of adaptive IRE1/XBP1s signaling. Consistent with this function, we find that pharmacologic inhibition of SLC33A1 with IXA4 selectively reduces viability of KEAP1-deficient lung adenocarcinoma cells that have elevated levels of glutathione, mimicking the sensitivity of these cells to genetic deletion of SLC33A1. Our work demonstrates a new physiologic role of SLC33A1 in regulation of ER redox homeostasis and designates IXA4 as a pharmacologic inhibitor of SLC33A1 that can be used to evaluate the biological impact and therapeutic utility of SLC33A1 inhibition in homeostasis and in disease.

## INTRODUCTION

SLC33A1 is an endoplasmic reticulum (ER) membrane transporter that has been reported to transport acetyl-CoA into the ER lumen for protein and ganglioside acetylation^1–3^. Alterations in SLC33A1 function have been implicated in the onset and pathogenesis of multiple etiologically distinct diseases. For instance, causal SLC33A1 mutations have been found in Huppke-Brendel syndrome and autosomal dominant spastic paraplegia^4–9^. Increased SLC33A1 activity has also been linked with development of various diseases, including autism spectrum disorder and intellectual disability^10^. More recently, SLC33A1 was identified as a vulnerability in loss-of-function KEAP1-mutant lung adenocarcinoma, where depletion of SLC33A1 blocked tumor cell proliferation and hindered tumorigenesis^11^. Haploinsufficiency of SLC33A1 has also been shown to enhance secretory proteostasis and reduce disease severity in mouse models of Alzheimer’s disease^12^. These findings pinpoint SLC33A1 as an attractive drug target to intervene in diverse diseases; however, no small-molecule modulators of SLC33A1 function have been described.

We previously characterized the small molecule IXA4 as a selective activator of the IRE1/XBP1s signaling arm of the Unfolded Protein Response (UPR)^13^. The UPR is the cellular mechanism that is activated to counter ER stress brought about by various anomalies, including protein misfolding and lipid imbalances^14,15^. UPR signaling, mediated by the ER stress sensors IRE1, ATF6, and PERK, acts initially in a protective manner to restore cellular homeostasis, inducing a transient attenuation of protein synthesis (downstream of PERK) and stimulating activation of the stress-responsive transcription factors XBP1s (downstream of IRE1), ATF6 (a cleaved form of full-length ATF6), and ATF4 (downstream of PERK)^14,15^. The activation of adaptive UPR signaling promotes remodeling of ER and cellular physiology to counter the initiating stress and foster cell survival in response to acute ER insults.^14,15^ However, if this protective response fails to rectify the imbalances, the UPR then activates inflammatory and apoptotic signaling pathways (chiefly due to PERK activity and facets of IRE1 signaling induced by chronic, severe ER stress) that lead to the cell’s demise^16^. The balance between protective and deleterious UPR signaling is often altered in disease, suggesting that pharmacologically increasing adaptive UPR signaling such as the IRE1/XBP1s pathway may be of therapeutic utility in diverse conditions^17,18^. Motivated by this notion, we set out to identify compounds that selectively increase adaptive IRE1/XBP1s signaling, resulting in the discovery of several selective activators of this protective UPR arm, including IXA4.^13^

IXA4 selectively induces adaptive – but not detrimental – IRE1 signaling in both cells and mice.^13,19^ Multiple groups, including us, have used this compound to study the functional role and therapeutic potential of pharmacologically increasing protective IRE1/XBP1s signaling in numerous cell-based and mouse models of disease. For instance, activation of IRE1/XBP1s signaling via treatment with IXA4 brings about beneficial remodeling of liver and pancreas in obese-diabetic mice and corrects multiple features of systemic metabolic dysfunction.^19^ Similarly, IXA4-induced IRE1/XBP1s activation boosts remyelination in primary cell models of Charcot-Marie-Tooth disease type 1B (CMT1B).^20^ In yet another example of the utility of this molecule, IXA4-induced IRE1/XBP1s signaling was shown to promote clearance of neurotoxic poly(GR) repeats in mouse models of amyotrophic lateral sclerosis (ALS).^21^ Nevertheless, despite its promise, the protein target and mechanism whereby IXA4 induces adaptive IRE1/XBP1s signaling was unknown, limiting its appeal for translational development.

To identify the protein target relevant for IXA4 action and uncover the molecular basis of its ability to selectively activate adaptive IRE1/XBP1s signaling, we applied complementary genetic and chemoproteomic strategies. We find that IXA4 binds the ER-localized membrane transporter SLC33A1 to induce IRE1/XBP1s signaling. A cryo-electron microscopy (cryo-EM) structure of IXA4 in complex with SLC33A1 shows that IXA4 binds the central channel of SLC33A1, occluding this channel and blocking metabolite transport. Though SLC33A1 was previously suggested as a transporter of acetyl-CoA into the ER for protein acetylation^1–3,22^, in our hands, neither genetic depletion nor IXA4-mediated inhibition of SLC33A1 altered acetyl-CoA trafficking into the ER or ER protein acetylation. Instead, we found that IXA4 treatment caused accumulation of oxidized glutathione in the ER, hyperoxidizing the ER and selectively inducing adaptive IRE1/XBP1s signaling. Our findings uncover a role for SLC33A1 in regulating ER reduction-oxidation (redox) balance and establish IXA4 as an inhibitor of SLC33A1 that can be used to study the physiologic outcome and therapeutic utility of targeting SLC33A1 in diverse settings/diseases.

## RESULTS

### SLC33A1 deletion selectively increases IRE1/XBP1s signaling and renders cells refractory to IXA4-induced IRE1 activation

To identify genes whose deletion disrupts IXA4-induced IRE1 activation, we generated a stable HEK293 cell line expressing Cas9 and an XBP1-Venus IRE1 reporter (**Fig. S1A**)^23^. Upon activation, IRE1 splices the mRNA encoding this reporter, resulting in a frameshift that allows expression of the fluorescent Venus protein. Flow cytometry analysis confirmed that treatment with IXA4 activates this reporter and that CRISPR-mediated deletion of *IRE1* blocked this activation (**Fig. 1A**). We used these cells in a genome-wide CRISPR screen to identify genes whose deletion blocked IXA4-induced IRE1 activation (**Fig. 1B**). We infected HEK293 cells expressing Cas9 and the XBP1-Venus reporter with a Brunello lentiviral library comprising 4 sgRNA guides per gene^24^. Infected cells were treated with vehicle or IXA4 for 14 h and sorted by Venus fluorescence, collecting the top and bottom 20% of fluorescent cells for each condition. Using next generation sequencing, we then identified sgRNAs enriched in these fractions and calculated gene scores for each perturbation with established computational pipelines.^25,26^ As expected, genetic depletion of ER proteostasis factors such as *BiP* and *MANF* increased XBP1-Venus reporter fluorescence both in the presence and absence of IXA4, confirming the effectiveness of the screen (**Fig. 1C**; **Table S1**). This screen identified *SLC33A1* as a gene whose deletion, on its own, increased XBP1-Venus fluorescence in vehicle-treated cells (**Fig. 1C, D**). This finding is consistent with other reports that found that deletion of *SLC33A1* activates IRE1/XBP1s signaling.^27,28^ Notably, treatment with IXA4 failed to further increase the XBP1-Venus signal in *SLC33A1-*deficient cells, indicating that loss of SLC33A1 induces IRE1 activity and renders mutant cells refractory to further IRE1 activation by IXA4.

**Figure 1.**
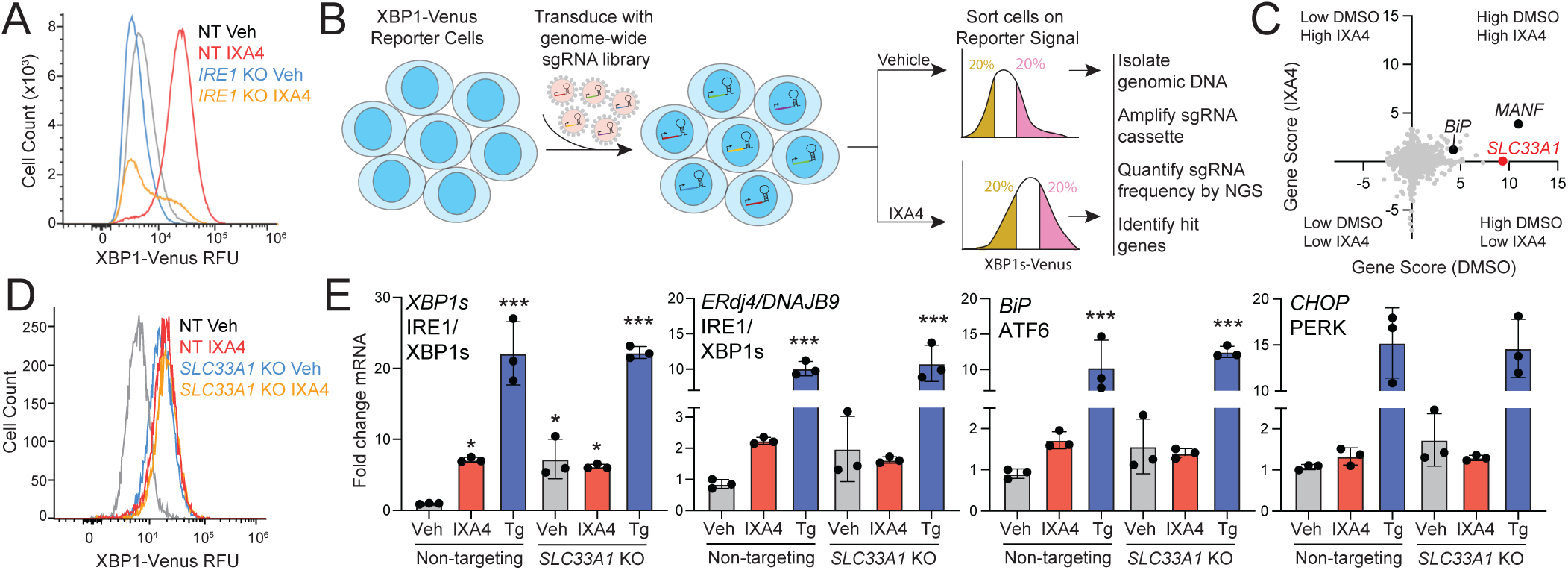
CRISPR screen identifies SLC33A1 as a protein involved in IXA4-induced IRE1/XBP1s signaling. **A**. XBP1-Venus reporter signal, measured by flow cytometry, in HEK293 cells stably expressing Cas9 and transduced with lentivirus encoding non-targeting (NT) or *IRE1* sgRNA treated for 14 h with vehicle or IXA4 (10 µM). **B**. CRISPR screen used to identify genes whose deletion blocks IXA4-induced IRE1 activation. **C**. Plot showing gene score (DMSO) vs. gene score (IXA4), quantified as previously defined^26^, from genome-wide CRISPR screen. **D**. XBP1-Venus signal, measured by flow cytometry, in HEK293 cells stably expressing Cas9 and transduced with lentivirus encoding non-targeting (NT) or *SLC33A1* sgRNA treated for 14 h with vehicle or IXA4 (10 µM). **E**. Expression, measured by qPCR, of the IRE1 target genes *XBP1s* and *DNAJB9*, the ATF6 target gene *BiP,* and the PERK target gene *CHOP* in HEK293 cells stably expressing Cas9 transduced with lentivirus encoding NT or *SLC33A1* sgRNA treated for 4 h with vehicle, IXA4 (10 µM) or Tg (0.5 µM). *p<0.05, ***p<0.005 one-way ANOVA.

To confirm these findings, we measured gene expression of *XBP1s* and the IRE1 target gene *DNAJB9/ERdj4* in HEK293 cells with CRISPR-mediated deletion of *SLC33A1* treated in the presence or absence of IXA4. *SLC33A1* deletion increased expression of *XBP1s* and *DNAJB9* to the same level seen in wild-type HEK293 cells treated with IXA4 (**Fig. 1E**). IXA4 treatment did not further increase expression of these genes in cells lacking *SLC33A1* (**Fig. 1E**). In contrast, treatment with the global ER stressor thapsigargin (Tg) induced robust expression of *XBP1s* and *DNAJB9* in both control and *SLC33A1-*deficient HEK293 cells. Deletion of *SLC33A1,* akin to IXA4 treatment, did not significantly increase expression of target genes regulated by ATF6 (e.g., *BiP*) or PERK (e.g., *CHOP*) (**Fig. 1E**), indicating that *SLC33A1* deletion shows the same preferential induction of IRE1/XBP1s signaling seen upon IXA4 treatment. Similar findings were made in HEK293 cells depleted of *SLC33A1* using shRNA and in MEF cells where *SLC33A1* was deleted using two different sgRNAs (**Fig. S1B, C**). Re-expression of SLC33A1 in *SLC33A1-*deficient HEK293 cells reduced *XBP1s* expression to basal levels and restored IXA4-induced increases in *XBP1s* expression, confirming that these effects are due to loss of SLC33A1 (**Fig. S1D**). *SLC33A1* deletion also induced IRE1-dependent increases in XBP1s protein and abolished any additional increase by IXA4, confirming increased IRE1/XBP1s signaling in *SLC33A1-*depleted cells at the protein level (**Fig. S1E**). These findings reveal SLC33A1 as a relevant protein that endows IXA4 with its ability to selectively activate IRE1/XBP1s signaling and suggests that IXA4 may act as a direct inhibitor of SLC33A1 function.

### IXA4 directly binds SLC33A1

To examine this possibility, we generated a fully functionalized derivative of IXA4, PTG2018 (**Fig. 2A**), bearing a diazirine moiety (to allow UV-light-mediated photocrosslinking and covalent capture of bound protein targets in live cells) and an alkyne handle (to enable conjugation of reporter tags via click chemistry for subsequent enrichment and identification of compound-bound proteins)^29–32^. PTG2018 increased activation of the IRE1-dependent XBP1-Renilla luciferase (RLuc) reporter in HEK293 cells, albeit with lower potency than that of IXA4 (**Fig. 2B**)^13^. Co-treatment with the IRE1 RNAse inhibitor 4µ8c blocked the increase in XBP1-RLuc signal induced by PTG2018, confirming that it was due to IRE1 activation (**Fig. S2A**)^33^. To assess the ability of PTG2018 to bind proteins in mammalian cells, we treated HEK293T cells with increasing doses of PTG2018, exposed them to UV light to crosslink PTG2018-bound proteins, prepared proteomes, and used click chemistry to append a tetramethylrhodamine (TAMRA) azide fluorophore to PTG2018-bound proteins prior to in-gel fluorescence analysis. This work showed that PTG2018 bound numerous proteins in a dose dependent manner (**Fig. S2B**). Next, to identify high occupancy targets of IXA4, we applied a chemoproteomic workflow using click-chemistry-mediated biotin conjugation, streptavidin affinity enrichment, and Tandem Mass Tag (TMT)-based quantitative proteomics to proteomes prepared from HEK293T cells treated with PTG2018 (5 µM) in competition with an excess of IXA4 (25 μM or 50 μM)^34^. This analysis identified three proteins for which IXA4 efficiently competed PTG2018 probe labeling at both IXA4 concentrations: SLC33A1, EPHX1, and SOAT1 (**Fig. 2C, D**; **Table S2**). In-gel analysis confirmed IXA4-dependent suppression of PTG2018 labeling of FLAG-tagged SLC33A1 and FLAG-tagged EPHX1 (**Fig. 2E** and **Fig. S2C**). Similarly, we noted IXA4-dependent competition of PTG2018 labeling of endogenous SOAT1 (**Fig. S2D**). However, unlike what was seen for *SLC33A1* (**Fig. 1C, D**), CRISPR-mediated deletion of either *EPHX1* or *SOAT1* did not alter activity of the XBP1-Venus reporter in HEK293 cells in the presence or absence of IXA4, indicating that IXA4 binding to these proteins does not affect IRE1 signaling (**Fig. S2E, F**). These observations show that SLC33A1 is the direct protein target of IXA4 that enables it to induce adaptive IRE1/XBP1s signaling.

**Figure 2.**
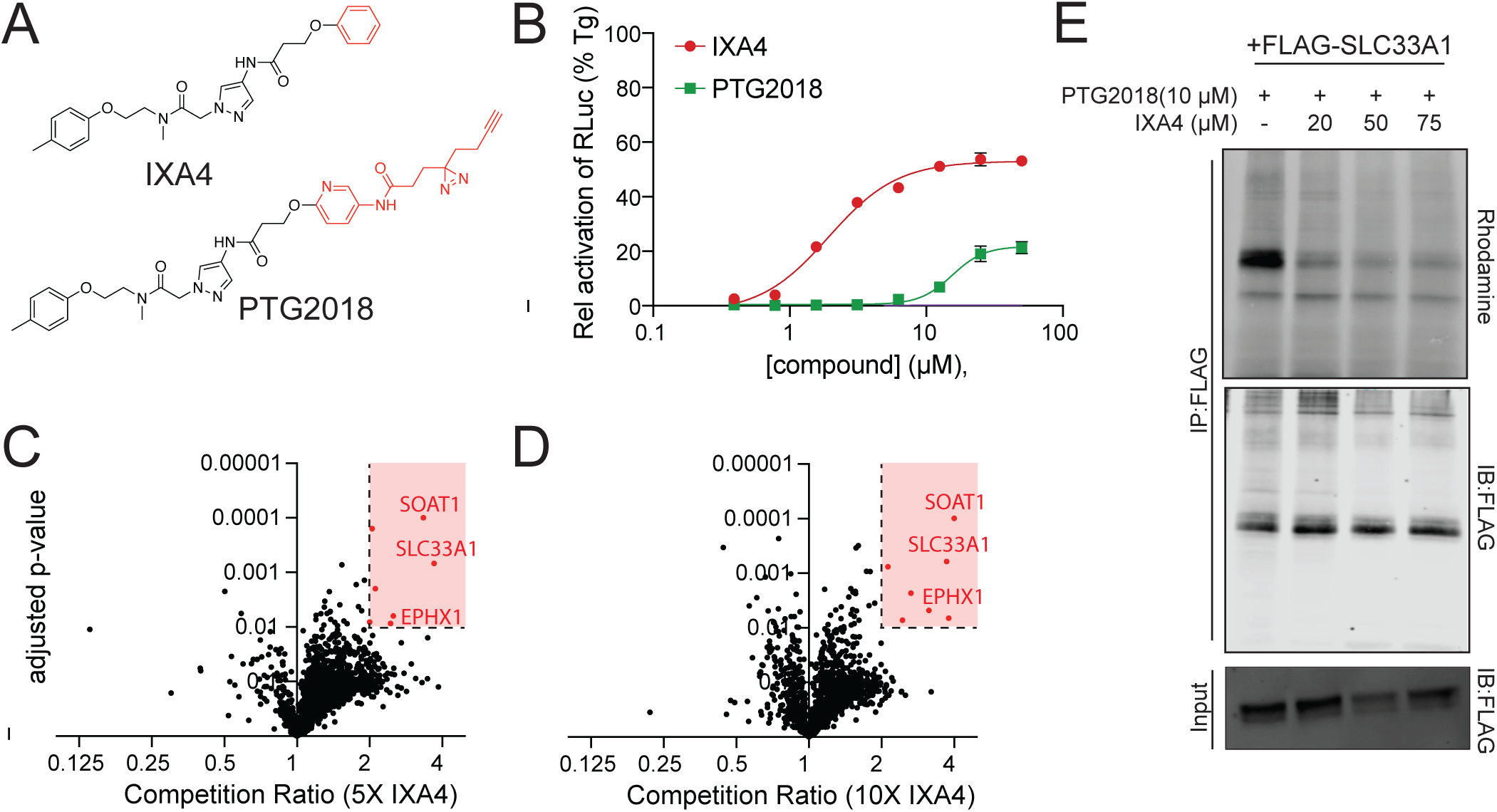
IXA4 binds SLC33A1. **A.** Structure of IXA4 and PTG2018. **B**. XBP1-RLuc signal, normalized to that observed in cells treated with thapsigargin (Tg), in HEK293 cells treated for 14 h with the indicated dose of IXA4 or PTG2018. **C**, **D**. Plots showing competition ratio versus adjusted *p*-value for proteins labeled in HEK293T cells treated for 30 min with PTG2018 (5 μM) in the presence of 5X IXA4 (25 μM; **C**) or 10X IXA4 (50 µM; **D**). **E**. In-gel fluorescence (top panel) and immunoblots of FLAG immunopurification and proteomes (input) from HEK293T cells overexpressing FLAG-tagged SLC33A1 and treated with the indicated concentration of PTG2018 with/without IXA4. PTG2018 labeling of FLAG-tagged SLC33A1 was visualized by appending TAMRA-azide to PTG2018 via click chemistry.

### IXA4 binds the central channel of SLC33A1

To understand the binding mode of IXA4 to SLC33A1, and its potential mechanism of action, we determined cryo-EM structures of full length SLC33A1 in the absence and presence of IXA4 at ∼3.3 Å and 3.27 Å resolution, respectively (**Fig. 3A, B**, **Fig. S3, S4, Table S3**). SLC33A1 adopts the canonical Major Facilitator Superfamily (MFS) fold, consisting of 12 transmembrane (TM) helices organized into two pseudo symmetric domains: TMs 1-6 (N-terminal domain, NTD) and TMs 7-12 (C-terminal domains, CTDs), with both the N- and C-termini facing the cytosol (**Fig. S5A**, **B**). A prominent loop between TM11 and TM12 extends into the ER lumen and forms a partially ordered soluble domain composed of two β-strands and a short α-helix stabilized by disulfide bonds (Cys479–Cys496 and Cys487–Cys503). The β-strands of this domain were sufficiently resolved for atomic modeling, but flexibility of the remainder of the luminal domain limited our ability to refine its atomic structure.

**Figure 3.**
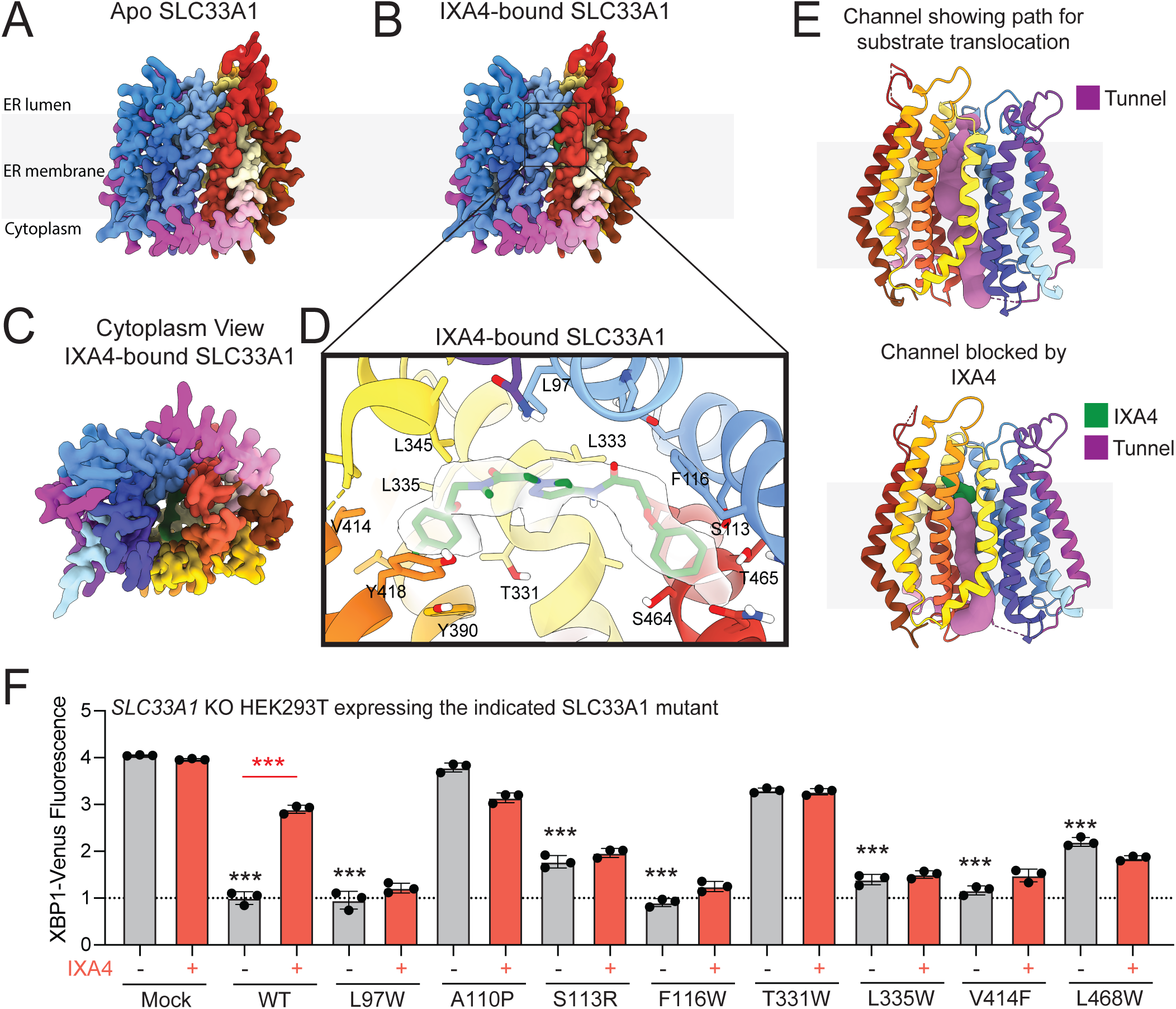
IXA4 binds the central channel of SLC33A1. **A**, **B**. CryoEM density maps of apo- and IXA4-bound SLC33A1**. C**. Cytoplasmic view of IXA4-bound SLC33A1. **D**. Close-up view highlighting the IXA4 binding pocket in SLC33A1. IXA4 is shown in green and the corresponding EM-density in semi-transparent surface; interacting residues are shown as sticks and labeled. **E**. Pathway analysis using MOLEonline shows a solvent accessible channel for apo (top) and IXA4-bound SLC33A1. **F**. XBP1-Venus signal, measured by flow cytometry, after reconstitution of wild-type SLC33A1 and mutant proteins in HEK293 XBP1-Venus reporter cells lacking endogenous SLC33A1 following treatment with vehicle or IXA4 (10 µM) for 14 h. ***p<0.005, one-way ANOVA; black asterisks represent comparisons between vehicle-treated cells expressing the indicated mutant and vehicle mock transfected cells. Red asterisks indicate comparisons between vehicle and IXA4-treated samples.

Our apo-SLC33A1 structure exhibits an ‘outward-open’ conformation, consistent with previously reported structures of full-length SLC33A1, wherein the central substrate binding channel is accessible from the cytosolic side (**Fig. 3C**)^28,35^. In this conformation, the N- and C-termini are splayed apart at the cytoplasmic interface between TM5 and TM8 forming a ∼50 Å wide cleft. The luminal gate, in contrast, is tightly sealed by close packing between TM1/2 and TM7/11, with additional contributions from TM5. A cytoplasmic lateral helix bridging TM6 and TM7 spans across the central cleft, capping the entrance and anchoring the two domains (**Fig. S5B**).

Our structure of IXA4-bound SLC33A1 closely resembles that of apo-SLC33A1 (**Fig. 3A-C**), although clear density for IXA4 could be seen within the central cavity at the NTD-CTD interface (**Fig. 3C, D**). Hydrophobic residues, including L333, L335, L345, and L97 stabilize one end of the IXA4 molecule. The aromatic part of IXA4 is stabilized by π-interactions with Y418 and Y390 at one side, while the opposite end is stabilized by an aromatic contact with F116 (**Fig. 3D**). In addition, IXA4 forms H-bond and electrostatic interactions with residues S113, T331, S464 and T465, further defining binding specificity and stability. Analysis of our SLC33A1 structures using MOLEonline^36^ uncovered a solvent accessible channel extending from the cytoplasmic face to the ligand binding pocket, suggesting a transport pathway through the ER membrane (**Fig. 3E**). The positioning of IXA4 within this channel suggests that compound binding occludes SLC33A1’s central channel, thereby blocking substrate transport.

To further characterize the interaction between IXA4 and SLC33A1, and its link to selective induction of adaptive IRE1/XBP1s signaling, SLC33A1 mutants predicted to disrupt IXA4 binding were expressed in *SLC33A1*-deficient HEK293 cells, and the activity of the XBP1-Venus IRE1 reporter was measured in the presence/absence of IXA4. This analysis identified multiple SLC33A1 mutants that, like wild-type SLC33A1, reduced the increased IRE1 activity seen in *SLC33A1*-null cells (e.g., L97W, F116W, L335W, V414F, and L468W; **Fig. 3F**). This observation indicates that these mutants retain normal SLC33A1 function. Interestingly, as predicted by our structural information, these mutants were all refractory to IXA4-induced increases in XBP1-Venus signal, denoting that they impair IXA4-mediated IRE1 activation. In contrast, re-expression of these mutants in SLC33A1-null cells did not impair IRE1 activation induced by the global ER stressor thapsigargin (Tg; **Fig. S5C**). Together with our structural observations, these findings support the notion that IXA4 selectively induces adaptive IRE1/XBP1s signaling by binding SLC33A1 and disrupting its transport of metabolites across the ER membrane.

### Inhibition of SLC33A1 hyperoxidizes the ER and activates IRE1 signaling

SLC33A1 has previously been suggested to transport acetyl-CoA into the ER to enable ER protein acetylation.^2,3^ However, we found that IXA4 treatment did not alter import of [^3^H] acetyl-CoA into ER microsomes isolated from treated HEK293T cells (**Fig. 4A**). ER microsomes isolated from *SLC33A1*-deficient HEK293T cells also did not show a reduction in [^3^H] acetyl-CoA import (**Fig. 4B**). Similarly, neither treatment with IXA4 nor *SLC33A1* depletion reduced acetyl-CoA levels in ER swiftly isolated from HEK293T cells (**Fig. 4C**). Moreover, we did not detect altered acetylation of ER proteins isolated from HEK293T cells treated with IXA4 or depleted of *SLC33A1*, as assessed using a chemical probe that reports on protein acetylation (**Fig. S6A, B**)^37^. These findings indicate that, in the contexts we have studied, pharmacologic inhibition or genetic deletion of SLC33A1 does not alter acetyl-CoA import into the ER or ER protein acetylation.

**Figure 4.**
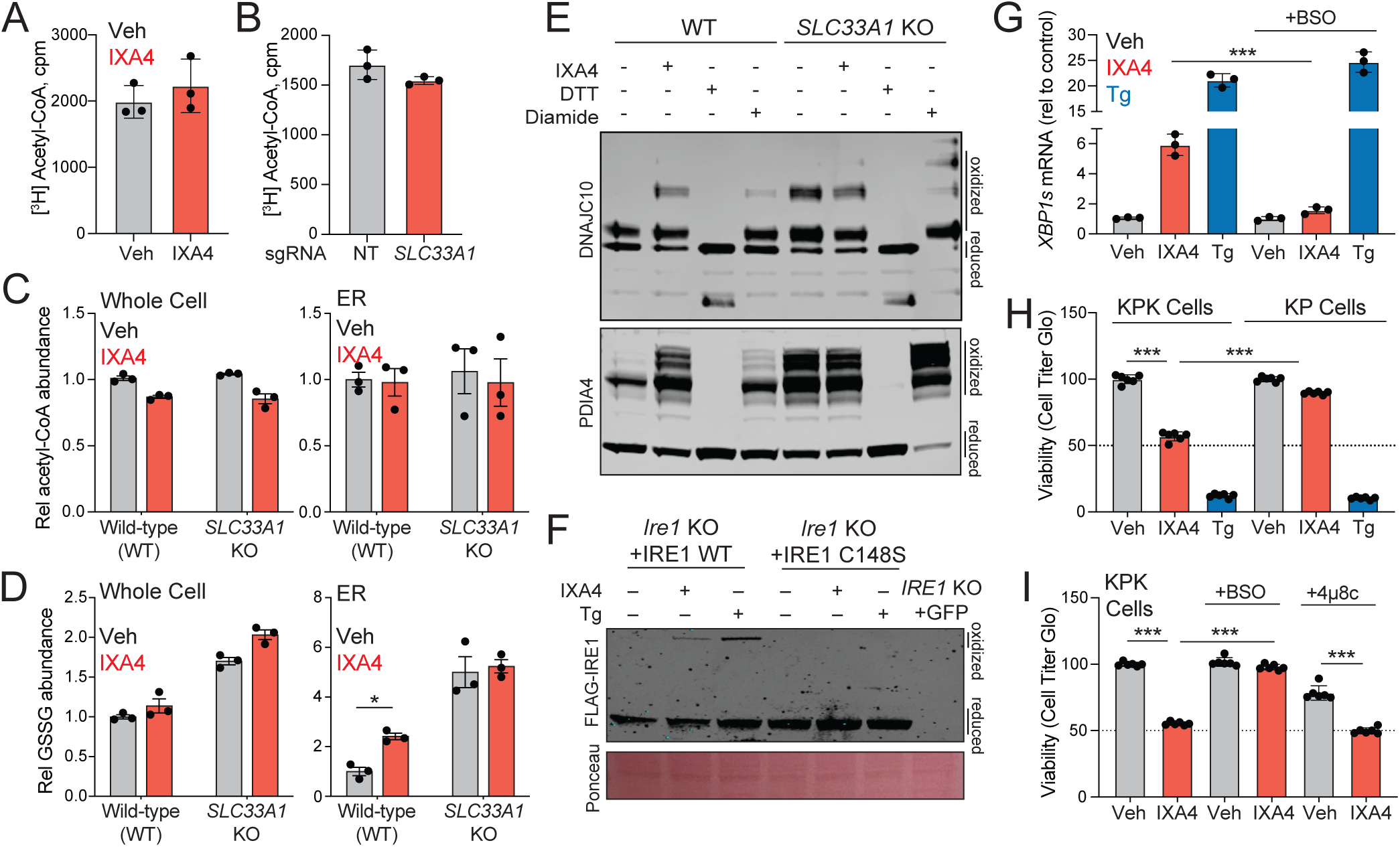
IXA4 binding to SLC33A1 promotes ER hyperoxidation and IRE1 activation. **A.** [^3^H]-acetyl-CoA uptake by ER microsomes (50 µg protein) isolated from HEK293T cells treated with/without IXA4 (10 µM) during the uptake assay. **B**. [^3^H]-acetyl-CoA uptake by ER microsomes (50 µg protein) isolated from HEK293T cells expressing Cas9 and non-targeting (NT) or *SLC33A1* sgRNA. **C**, **D**. Relative acetyl-CoA abundance (**C**) and oxidized glutathione abundance (GSSG; **D**) in whole cell or ER isolated from HEK293T cells with non-targeting (NT) sgRNA or sgRNA targeting SLC33A1. Cells were treated with IXA4 (10 µM) for 4 h before ER isolation and LC/MS analysis. **E**. Immunoblot of DNAJC10 and PDIA4 in proteomes prepared from HeLa cells treated for 1 h with IXA4 (10 µM), DTT (5 mM), or diamide (500 µM). Following lysis in the presence of alkylating agents, disulfides within these proteins were reduced and then labeled with PEG(5K)-maleimide (2 mM), allowing visualization of oxidized and reduced protein populations. **F**. Immunoblot of proteomes from *Ire1^-/-^* MEFs reconstituted with wild-type or a C148S IRE1 mutant, treated for 2 h with IXA4 (10 µM) or thapsigargin (Tg; 0.5 µM), and separated on a non-reducing SDS-PAGE gel. Oxidized and reduced IRE1 populations are shown. **G**. *XBP1s* levels, measured by qPCR, in HEK293T cells pre-treated for 18 h with BSO (50 µM) and then treated for 3 h with IXA4 (10 µM). **H**. Viability, assessed with CellTiter-Glo, of KP and KPK cells treated for 3 days with IXA4 (10 µM) or thapsigargin (Tg; 0.5 µM). **I**. Viability, assessed with CellTiter-Glo, of KP and KPK cells treated for 3 days with IXA4 (10 µM), BSO (50 µM), and 4µ8c (32 µM). *p<0.05, ***p<0.005, one-way ANOVA.

More recently, SLC33A1 was shown to regulate ER redox balance by mediating export of oxidized glutathione (GSSG) from the ER to the cytosol^28^, raising the possibility that pharmacological inhibition of SLC33A1 would hyperoxidize the ER lumen. Published DNA microarray data show that ER hyperoxidation brought about by overexpression of the ER oxidase ERO1 preferentially induces IRE1/XBP1s target genes, relative to PERK targets (**Fig. S6C**)^38^. This observation hinted that ER hyperoxidation engendered by IXA4-mediated SLC33A1 inhibition may explain the selective induction of IRE1/XBP1s signaling seen with this compound. In line with this notion, treatment of HEK293T cells with IXA4 increased ER levels of GSSG, without altering total cellular GSSG levels (**Fig. 4D**). A similar increase in ER GSSG was seen in *SLC33A1-*deficient HEK293T cells (**Fig. 4D**). Moreover, akin to what was noted in *SLC33A1*-null cells^28^, treatment with IXA4 increased the oxidation status of the ER-localized protein disulfide isomerases PDIA4 and DNAJC10 (**Fig. 4E**). These data show that, by inhibiting SLC33A1, IXA4 increases ER GSSG levels, hyperoxidizing the ER lumen.

IRE1 activity is known to be affected by redox-dependent regulation of cysteines residues in its luminal domain^39^. Thus, we considered the prospect that hyperoxidation of the ER lumen triggered by IXA4-dependent inhibition of SLC33A1 may induce IRE1 signaling via redox-dependent modification of IRE1’s luminal cysteines. In support of this possibility, we noted that IXA4-treatment increased disulfide-linked IRE1 oligomers mediated by C148 of the IRE1 luminal domain (**Fig. 4F**). However, mutation to serine of all three cysteine residues within the IRE1 luminal domain did not impair the ability of IXA4 to activate IRE1 and induce splicing of *XBP1* (**Fig. S6D**). This observation indicates that IXA4-induced activation of IRE1 signaling does not depend on redox regulation of its luminal cysteine residues. Nonetheless, treatment of HEK293T cells with BSO (L-buthionine sulfoximine), an inhibitor of glutamate cysteine ligase catalytic (GCLC) subunit, a key enzyme in GSH synthesis, completely abolished the ability of IXA4, but not thapsigargin, to induce IRE1/XBP1s signaling (**Fig. 4G).** Because BSO limits cellular glutathione production, thereby reducing the pool of GSH available for oxidation within the ER, this result is consistent with a model in which IXA4-induced IRE1/XBP1s activation relies on ER GSSG accumulation. Together, our data show that IXA4-mediated inhibition of SLC33A1 selectively activates IRE1/XBP1s signaling via a mechanism that involves accumulation of GSSG in the ER and subsequent hyperoxidation of the ER lumen.

### Pharmacologic SLC33A1 inhibition preferentially decreases viability of Keap-1 deficient lung adenocarcinoma cells

SLC33A1 has been identified as a vulnerability of *Kras^G12D/+^*; *p53^−/−^*; *Keap1^−/−^* (KPK) lung adenocarcinoma cells^11^. Because they lack *Keap1*, the cell’s primary sensor for oxidative stress, KPK cells hyperactivate the antioxidant program regulated by nuclear factor, erythroid 2-like 2 (NFE2L2 or NRF2)^40^. Genetic depletion of *SLC33A1* reduces viability of KPK cells, but not parental *Kras^G12D/+^*; *p53^−/−^* (KP) cells. The selective sensitivity of KPK cells to *SLC33A1* deletion is attributed to both, increased GSH biosynthesis resulting from NRF2 hyperactivation, and the reduction in SLC33A1-mediated export of GSSG from the ER^11,28^. We tested the extent to which, akin to genetic deletion^11^, IXA4-mediated pharmacological inhibition of SLC33A1 would preferentially reduce KPK cell growth, relative to KP cells. Treatment with IXA4 did not impact viability of KP cells but significantly reduced survival of KPK cells (**Fig. 4H**, **Fig. S6E**). Blocking GSH synthesis in KPK cells using BSO abolished IXA4’s ability to reduce KPK cell viability **(Fig. 4I**, **Fig. S6F**), mirroring results seen with *SLC33A1* deletion^11^. However, co-treatment of cells with IXA4 and the IRE1 RNAse inhibitor 4µ8c did not blunt IXA4’s effect on KPK cell proliferation (**Fig. 4I)**. These observations indicate that pharmacologic inhibition of SLC33A1 selectively reduces KPK cell viability in a manner dependent on dysregulation of glutathione levels, but not on increased IRE1/XBP1s signaling.

## DISCUSSION

Pharmacological activation of adaptive signaling through the IRE1/XBP1s arm of the UPR using compounds like IXA4 has emerged as a potential therapeutic strategy to intervene in multiple conditions, including metabolic and neurodegenerative disorders^17–21^. Here, we applied a multipronged strategy to identify SLC33A1 as the protein target of IXA4 responsible for its selective activation of IRE1/XBP1s signaling. IXA4 binds in the central cavity of SLC33A1, occluding its ability to transport metabolites across the ER membrane. Notably, we find that pharmacologic inhibition or genetic depletion of SLC33A1 increases the levels of oxidized glutathione within the ER, hyperoxidizing the ER lumen to induce activation of adaptive IRE1/XBP1s signaling. Moreover, mirroring results seen with genetic deletion of SLC33A1^11^, IXA4-mediated inhibition of SLC33A1 selectively reduced proliferation of KEAP1-mutant lung adenocarcinoma cells that have increased glutathione levels, a finding consistent with its mechanism of action involving alterations in ER redox balance. Beyond our identification of IXA4 as the first described pharmacologic inhibitor of SLC33A1 and a useful tool to study the role of SLC33A1 in physiology and disease, our findings support a new function for SLC33A1 as a key regulator of ER redox homeostasis^28^. Furthermore, they reveal an adaptive mechanism, IRE1/XBP1s signaling, triggered to correct ER redox imbalances.

SLC33A1 has been thought of as an ER acetyl-CoA importer required for acetylation of ER resident proteins^2,3,22^. While this may be an important function of SLC33A1 in certain cell types, in the systems we have studied neither pharmacologic inhibition of SLC33A1 with IXA4 nor genetic depletion of *SLC33A1* altered acetyl-CoA import into isolated ER, ER acetyl-CoA levels, or ER protein acetylation. These data indicate that SLC33A1 must have additional/alternative functions beyond acetyl-CoA trafficking. Consistent with this notion, a recent report elegantly demonstrated that deletion of *SLC33A1* blocks the export of oxidized glutathione (GSSG) from the ER, increasing ER levels of GSSG and hyperoxidizing the ER environment^28^. We too found that pharmacologic or genetic inhibition of SLC33A1 increases ER GSSG levels and hyperoxidizes the ER, as reflected in greater oxidation of protein disulfide isomerases within the ER lumen. These findings establish a new role for SLC33A1 in regulating ER redox homeostasis through the export of GSSG – a critical activity as no glutathione reductase is present in the mammalian ER lumen^41^. The critical role of SLC33A1 in redox homeostasis is also supported by multiple CRISPR screens that identified SLC33A1 as an important regulator of cellular sensitivity to genetic (e.g., KEAP1 deficiency)^11^ or chemical (e.g., arsenic, cadmium) oxidative insults^42^.

Maintaining a proper oxidized/reduced glutathione (GSSG/GSH) balance is critical for oxidative protein folding within the ER, the process of disulfide bond formation in nascent proteins and the correction of improper disulfide bonds in misfolded proteins^43–45^. Hence, it is perhaps intuitive that alterations in the ER GSSG/GSH ratio, as we saw with inhibition or deletion of SLC33A1, would activate adaptive UPR signaling to restore homeostasis. Indeed, other studies have noted an activation of UPR signaling upon loss of SLC33A1 function^27,28,46^. Prior work has also shown that ER hyperoxidation brought about by ERO1 overexpression can preferentially activate the IRE1/XBP1s, relative to PERK, arm of the UPR^38^. Moreover, XBP1s regulates the expression of many ER redox genes, including *PDIA4*, *DNAJC10*, and *ERO1B*, supporting a model whereby activation of IRE1/XBP1s signaling induced by SLC33A1 inhibition fosters restoration of ER redox homeostasis following acute redox stress^47,48^.

Nonetheless, because IXA4 is a selective activator of IRE1/XBP1s signaling, the question remains as to how pharmacological inhibition of SLC33A1 exclusively activates this adaptive UPR pathway. It may be that IRE1 is more responsive to ER hyperoxidation than other ER stress sensors. While we found that activation of IRE1 signaling due to pharmacologic SLC33A1 inhibition is independent of disulfide sensing within the IRE1 luminal domain, as mutation of all three Cys residues within this domain did not block IXA4-induced IRE1 activation, it may be that IXA4-induced ER hyperoxidation leads to broader changes in ER proteostasis that are preferentially sensed by the IRE1 luminal domain. For instance, hyperoxidation of multiple ER-resident PDIs in IXA4-treated cells, as we saw for PDIA4 and DNAJC10, may integrate to transiently activate IRE1 to promote downstream XBP1s signaling. We posit that this putative increased sensitivity of IRE1 to redox imbalances would allow the cell to trigger an adaptive response to oxidative stress without engaging the apoptotic machinery associated with global, chronic UPR activation. Consistent with this model, IXA4 activates IRE1/XBP1s signaling to more modest levels than global UPR activating compounds (e.g., Tg), potentially reflecting the transient nature of IRE1 activation afforded by the ER hyperoxidation induced by this compound.^13^ Ultimately, the finding that SLC33A1 is the relevant target for IXA4’s effects reveals new insight into the functional relationship between ER redox homeostasis and UPR pathway activation that can be harnessed to selectively activate adaptive IRE1/XBP1s signaling.

Identification of SLC33A1 as the protein target of IXA4 highlights the potential utility of pharmacologically targeting this transporter in multiple diseases, which our structural and mechanistic framework now enables. IXA4-induced activation of IRE1/XBP1s signaling has proven beneficial in cell-based and *in vivo* models of varied diseases, including obesity-diabetes, CMT1B, and ALS^13,19–21^. Pharmacological SLC33A1 inhibition has also been proposed as an attractive strategy to target KEAP1-mutant lung adenocarcinoma^11^. Our data support this suggestion, as IXA4 treatment selectively decreased viability of KEAP1-null lung adenocarcinoma cells. Finally, the importance of SLC33A1 in export of oxidized glutathione from the ER of oxidized glutathione to maintain ER redox balance offers a new context to reevaluate the molecular basis of disease-linked *SLC33A1* mutations in diseases such as Huppke-Brendel syndrome and autosomal dominant spastic paraplegia^4^.

## MATERIALS AND METHODS

### Chemicals and Reagents

IXA4 (WuXi AppTec), thapsigargin (Fisher Scientific, Cat # 50-464-295), l-buthionine sulfoximine (BSO) (Selleckchem, Cat # S9728), 4µ8c (MedChemExpress, Cat # HY-19707), diamide (Sigma, Cat # D3648), dithiothreitol (DTT) (Sigma, Cat # D3648-1G), puromycin (Gibco, Cat # A1113803), blasticidin (Gibco, Cat # A1113903), (^3^H)-acetyl-CoA (Revvity, Cat # NET290050UC), 3-butynoic acid (MedChemExpress, Cat # HY-W004784), N-ethylmaleimide (NEM) (Thermo Fisher Scientific, Cat # 23030), PTG2018 (WuXi AppTec), Q5 Site-Directed Mutagenesis Kit (New England Biolabs, E0554), NEBuilder HiFi DNA Assembly Master Mix (New England Biolabs, E2621), Quick-DNA Midiprep Plus Kit (Zymo Research, D4075), Quick-RNA Mini Kit (Zymo Research), High-Capacity cDNA Reverse Transcription Kit (Applied Biosystems, 4368814), Power SYBR Green PCR Master Mix (Applied Biosystems, 4367659), Endoplasmic Reticulum Isolation Kit (Millipore Sigma, ER0100), CellTiter-Glo Luminescent Cell Viability Assay (Promega, Cat # G7572), DC Protein Assay (Bio-Rad), Dulbecco’s Modified Eagle’s Medium (DMEM; Corning), Roswell Park Memorial Institute 1640 Medium (RPMI 1640; Corning), Fetal Bovine Serum (FBS) (Omega Scientific), L-Glutamine (Gibco), Penicillin-Streptomycin (Gibco), Dulbecco’s Phosphate-Buffered Saline (DPBS; Gibco), TrypLE Express Enzyme (Thermo Fisher Scientific), TransIT-Lenti Transfection Reagent (Mirus Bio, MIR 6600), Crystal Violet, Sequencing Grade Modified Trypsin (Promega, V5111), TMT Labeling Reagents (Thermo Fisher Scientific), Pierce High pH Reversed-Phase Fractionation Kit (Thermo Fisher Scientific, 84868), Streptavidin-Agarose Beads (Thermo Fisher Scientific, 20349), Biotin-PEG3-azide (AZ104), PEG(5k)-maleimide (Sigma, Cat # JKA5036-1G), and tetramethylrhodamine (TAMRA) azide (synthesized in house). Antibodies used include FLAG (Cell Signaling, Cat # 8146), DNAJC10 (Proteintech, Cat # 13101-1-AP), PDIA4 (Proteintech, Cat # 14712-1-AP), IRE1alpha (Cell Signaling, Cat # 3294), XBP1s (Cell Signaling, Cat # 40435), BiP (Cell Signaling, Cat # 3177), CHOP (Cell Signaling, Cat # 2895), and Tubulin (Cell Signaling, Cat # 2144).

### DNA constructs

Sequences of oligonucleotides used for cloning are provided in the **Table S4**. The XBP1s-Venus reporter was generated by replacing the Renilla in our published XBP1s-Renilla vector with Venus through NEBuilder (NEB, E2621)^23^. To clone individual sgRNAs, top and bottom oligonucleotides (IDT) were annealed and ligated to lentiviral sgRNA expression vectors LentiGuide-puromycin (Addgene #52963), LentiCRISPRv2-puromycin (Addgene #98290), or LentiCRISPRv2-blasticidin (Addgene #98293)). Four-sgRNAs expression constructs were generated using the pYJA5 (Addgene #217778) vector with an optimized protocol of Golden Gate assembly. Protospacer sequences for sgRNAs are listed in the **Table S4**. Point mutants of overexpressing plasmids (IRE1 and SLC33A1) were generated using the Q5 Site-Directed Mutagenesis Kit (New England Biolabs, E0554) following the manufacturer’s instructions. Primers and T_­_temperature were designed using the NEBaseChanger online tool. Plasmids were amplified in DH5α cells and isolated using the ZymoPURE Plasmid Miniprep Kit. All plasmid sequences were confirmed (Plasmidsaurus) with custom analysis and annotation.

### Cell culture

HEK293, HEK293T, and HeLa cells were purchased from ATCC. WT and IRE1^-/-^ MEF cell lines were a gift of David Ron. KP cells were provided by Tyler Jacks. HEK293, HEK293T, HeLa, and MEF cells were cultured in high-glucose Dulbecco’s Modified Eagle’s Medium (DMEM) (Corning) supplemented with 10% fetal bovine serum (FBS) (Omega Scientific), 2 mM L-glutamine (Gibco), 100 U/mL penicillin and 100 µg/mL streptomycin (Gibco). KP and KPK cells were cultured in RPMI 1640 (Corning) with 10% fetal bovine serum (FBS) (Omega Scientific), 2 mM L-glutamine (Gibco), 100 U/mL penicillin and 100 µg/mL streptomycin (Gibco). Cells were grown in standard culture conditions (37°C, 5% CO_­_) and tested for mycoplasma every 6 months. HEK293 Cas9 and KP Cas9 cells were generated via lentiviral infection with lentiCas9-Blast (Addgene #52962). dCas9-Zim3-KRAB HEK293 and HeLa cells were generated by lentiviral infection with pHR-UCOE-SFFV-Zim3-dCas9-P2A-Hygro (Addgene #188768). The HEK293 Cas9 cell line was further used to establish a reporter cell line by lentiviral infection with a XBP1s-Venus reporter vector. FACS-based sorting was used to isolate a clonal Cas9-XBP1s-Venus-HEK293 cell line for the CRISPR screen.

### Genome-wide CRISPR screen

To obtain pooled sgRNA viruses, a Brunello sgRNA library (Addgene # 73178) was transfected into HEK293T cells together with lentiviral plasmid packaging mix, pMD2.G (Addgene #12259) and psPAX2 (Addgene #12260), using TransIT-Lenti Transfection Reagent (Mirus, MIR 6600). The viral library was titrated to achieve a multiplicity of infection ∼0.3. Cas9-XBP1s-Venus-HEK293 cells were infected with the pooled sgRNA viral library, aiming for 1,000-fold representation of each guide, and selected with puromycin (2.5 µg/ml) for 3 days. Cells were then cultured for 2 days in the absence of puromycin. Cell populations were cultured such that a representation of at least 1,000 cells per sgRNA was maintained throughout the screen. Cells were seeded at 2.5 million cells per 10 cm dish (12 dishes total) in a volume of 7.5 mL media on day 0. On day 1, an additional 2.5 mL of media with DMSO or IXA4 (10 µM final concentration) were added into each dish. Following 16 h of DMSO or IXA4 treatment, cells were FACS-sorted and the cell populations in the top 20% and bottom 20% of Venus signal collected. Genomic DNA was extracted using Quick-DNA Midiprep Plus Kit (Zymo Research, catalog # D4075). Illumina sequencing libraries were prepared by first amplifying the sgRNA-containing vector sequences from the gDNA and then using a second PCR reaction to append sequencing adapters^49^. Libraries were sequenced on an Illumina NextSeq 500 (single end 75 bp reads). The MaGeCK count function was used to align reads to the Brunello sgRNA library and generate read count tables for all samples^25^. The phenotype and *p* value for each gene were calculated using an established bioinformatics pipeline^26^. Gene scores were defined as the product between the phenotype (logFC) and −log10(P value).

### In situ labeling of cells by IXA4

HEK293T cells grown in 10 cm plates were incubated in serum-free DMEM with the indicated probe and IXA4 competitor concentration or vehicle for 30 min at 37°C under a 5% CO_­_atmosphere. Treated cells were then irradiated with UV light (365 nm, Stratagene, UV Stratalinker 1800) for 20 min, harvested, washed twice with ice-cold DPBS (1,400*g*, 4°C, 3 min), and collected into 15 mL Falcon tubes. Cell pellets were resuspended in ice cold DPBS with 1X protease inhibitor and lysed by sonication (Branson probe sonifier; on: 15 ms; off: 40 ms, 15% amplitude). Protein concentrations were normalized (1 mg/mL, 50 μL final volume) using the Lowry protein assay. 6 μL of freshly prepared ‘click’ mixture (3 μL of 1.7 mM TBTA in 4:1 *t*-BuOH:DMSO, 1 μL of freshly-prepared 50 mM TCEP in DPBS, 1 μL of 1.25 mM tetramethylrhodamine (TAMRA) azide in DMSO and 1 μL of 50 mM CuSO_­_in water, added in that order) were added to each sample and the reaction allowed to proceed at room temperature for 1 h. The reaction was quenched by adding 17 μL of 4X SDS gel loading buffer followed by analysis of 50 μL of each reaction on a 10% acrylamide SDS-PAGE gel. Gels were imaged using Bio-Rad ChemiDoc MP Imagining in-gel fluorescence.

### Preparation of samples for mass spectrometry (MS) analysis

Cell lysates were prepared as stated above. To 0.5 mL of 1 mg/mL protein sample, 55 μL of freshly prepared ‘click’ mixture (30 μL of 1.7 mM TBTA in 4:1 *t*-BuOH:DMSO, 10 μL of freshly-prepared 50 mM TCEP in DPBS, 5 μL of biotin-PEG4-azide and 1 μL of 50 mM CuSO_­_in water, added in that order) were added and the reactions incubated at room temperature for 1 h. Samples were then precipitated with 3 mL cold methanol at -20°C for 1 h. The solution was centrifuged at 5000*g* for 10 min at 4°C to obtain protein pellets. The supernatant was discarded and the pellet was washed twice with 4:1 MeOH: CHCl_­_solution. The resulting pellets were solubilized in 500 μL of proteomics grade 6M urea in DPBS, containing 10% SDS by sonication. This was followed by addition of 50 μL of 1:1 mixture of freshly prepared TCEP (200 mM in DPBS) and K_­_CO_­_(600 mM in DPBS) and incubation at 37°C for 30 min. After reduction, the samples were alkylated by addition of 70 μL of 400 mM iodoacetamide in DPBS in room temperature for 30 min. The reaction was quenched by adding 130 μL of 10% SDS in DPBS and then diluted to 5.5 mL with DPBS. Next, 100 μL of pre-equilibrated streptavidin-agarose bead suspension (Thermo Fisher Scientific, 20349) was added to each sample for probe-labeled protein enrichment. The samples were rotated at room temperature for 1.5 h, centrifuged at 2000 rpm for 2 min and washed sequentially with 0.2% SDS in DPBS (5 mL), DPBS (5 mL), water (5 mL), and 100 mM TEAB pH 8.5 (5 mL). The beads were then transferred to 1.5 mL LoBind microcentrifuge tubes and digested overnight at 37°C in trypsin (Promega, V5111) premix in TEAB buffer (2 mL, 100 mM, pH 8.5) with 20 μL 100 mM CaCl_­_. The beads were centrifuged at 2000 rpm for 5 min, the supernatant was transferred to a new tube, and the beads were further washed with 100 μL TEAB buffer. The digest was labeled with TMT (Thermo Fisher Scientific) tags, incubated at room temperature for 1 h. The reaction was quenched by adding 6 μL of 5% hydroxylamine in water and incubated for 15 min, followed by drying under vacuum centrifugation at 45°C, 10 psi. Samples were then combined and fractionated using a Pierce High pH Reversed-Phase Fractionation Kit (Thermo Fisher Scientific, 84868), according to the manufacturer’s instructions, into six final fractions and dried and stored at -80°C until ready for MS analysis.

### Liquid chromatography–mass spectrometry analysis of TMT samples

TMT-labeled peptides were resuspended in MS sample buffer (0.1% formic acid in water) before being subjected to liquid chromatography-mass spectrometry (LC-MS) analysis as previously described^34^. In general, samples were loaded and eluted using an UltiMate 3000 RSLCnano system (Thermo Fisher Scientific) with a 220-min gradient separation method. The eluents were then analyzed and quantified in an Orbitrap Fusion Lumos mass spectrometer (Thermo Fisher Scientific). MS1 spectra were specified in 375–1,500-m/z scan range with a resolution of 120,000; peptides isolated for MS2 spectra were fragmented by collision-induced dissociation (CID) (30% collision energy); and synchronous precursor selection^50^ was used to isolate up to 10 MS2 ions for the MS3-based quantification through high-energy collision-induced dissociation (HCD) (65% collision energy).

### Real-time quantitative PCR (RT-qPCR) and RNAseq

Total RNA was isolated using the QuickRNA mini kit (Zymo) according to the manufacturer′s instructions. cDNA was synthesized from 1000 ng RNA using the High-Capacity cDNA Reverse Transcription Kit (Applied Biosystems, 4368814). RT-qPCR was performed using the Power SYBR Green PCR Master Mix (Applied Biosystems, 4367659), with data analyzed using the 2−^ΔΔC^t method and normalized to 36B4 expression. Primers were purchased from IDT; sequences used are listed in **Table S4**.

### Western blot analysis

Samples separated by SDS–PAGE were transferred onto nitrocellulose membranes. Membranes were incubated in blocking buffer (fat-free goat milk 5% w/v in Tris-buffered saline [TBS]) for 1 h at room temperature. Membranes were incubated overnight at 4°C with primary antibodies diluted in Bovine Serum Albumin (BSA)-containing TBS buffer overnight, washed three times for 15 min with TBS-Tween 0.1%, and incubated for 1 h at room temperature with fluorophore-conjugated secondary antibodies diluted in blocking buffer (1:20,000 dilution). The antibodies used in this work are listed above and all were used all used.at 1:1000 dilution.

### ER isolation, acetyl-CoA import, and protein acetylation assays

ER isolation was performed using a Millipore Sigma kit (ER0100), following the manufacturer’s protocol. Two 15 cm dishes of HEK293T cells were used per condition. 50 µg of ER isolated from non-targeting control cells and *SLC33A1* KO cells was exposed to 10 µM [^3^H] acetyl-CoA for 5 min. Treated ER was pelleted (100,000*g*, 30 min), washed with DPBS twice and resuspended in liquid scintillation solvent. In the case of *in vitro* IXA4 treatment, ER isolated from HEK293T cells was treated with IXA4 (10 µM) or DMSO for 15 or 30 min. [^3^H] acetyl-CoA at 10 µM was then added for 5 min. The treated ER was pelleted (100,000*g*, 30 min), washed twice with DPBS and resuspended in liquid scintillation solvent. Measurements were performed using a Beckman Coulter LS6500 Liquid Scintillation Counter. To monitor protein acetylation, HEK293T cells were treated with IXA4 (10 µM) for 3 h, followed by 1 h treatment with 3-butynoic acid (10 mM)^37^. Similarly, HEK293T cells bearing Cas9 with non-targeting or *SLC33A1* sgRNAs were treated with 3-butynoic acid (10 mM) for 1 h. ER microsomes were isolated and lysed by sonication in cold DPBS buffer. Lysates were normalized to 1 mg/mL in 44 µL and an azide-alkyne cycloaddition reaction (i.e., click chemistry) was performed by adding 6 µL of click-master mix (1 µL 1.25 mM TAMRA-azide, 1 µL 50 mM CuSO4, 3 µL 1.7 mM TBTA in 1:4 DMSO: *t*-butanol, 1 µL 50 mM TCEP) and incubation for 1 h at room temperature. Samples were resolved by SDS-PAGE and in-gel fluorescence acquired using a Bio-Rad Chemidoc imager.

### Flow cytometry analysis of XBP1s-Venus reporter cells

Reporter cells were seeded at a density of 20,000 cells per well in 96-well TC-treated flat bottom plates. The following day, cells were treated for 16 h with the indicated compound concentration. Following treatment, cells were dissociated using TrypLE Express (ThermoFisher). The enzymatic reaction was neutralized by addition of flow buffer (1% FBS, 1% Pen-Strep, 25 mM HEPES, 2.5 mM EDTA in DPBS). Flow cytometry was performed on a NovoCyte 3000 with NovoSampler Pro. Venus (568/592 nm) signal was measured with a FITC filter (530/30) and analyzed using FlowJo Software (BD Biosciences).

### XBP1 splicing assay

cDNA synthesis was performed from 1 µg of total RNA extracted from treated cells. *Xbp1s* and *Xbp1u* were amplified by using primers provided in **Table S4**. Products were resolved on a 5% agarose gel.

### Protein oxidation assay

To monitor the oxidation status of ER-resident proteins, a PEG-switch assay was performed as previously described^51^. Cells were treated with IXA4 (10 µM), DTT (5 mM), or diamide (500 µM) for 1 h. Following treatment, cells were washed with ice-cold DPBS containing 100 mM N-ethylmaleimide (NEM) to block free thiols and prevent post-lysis oxidation. Cells were lysed in lysis buffer (100 mM Tris pH 7.4, 1% SDS and 100 mM NEM) by sonication. Lysates were incubated at 50°C for 25 min to ensure complete alkylation of free cysteines. Excess NEM was removed using Zeba spin desalting columns (Pierce). 50 mM dithiothreitol (DTT) was added to each sample and incubated for 20 min at room temperature to reduce reversibly oxidized cysteine thiols. Excess DTT was removed using Zeba spin desalting columns. Reduced desalted samples were labeled with 2 mM PEG5000-maleimide (Sigma) in 0.5% SDS for 2 h at room temperature. The reaction was quenched by the addition of SDS-PAGE sample buffer. Samples were resolved by SDS-PAGE and analyzed by immunoblotting for DNAJC10 and PDIA4. The oxidized fraction is visualized as a higher molecular weight band due to the conjugation of the 5 kDa PEG moiety.

### Cell proliferation assays

Cell viability was assessed using the CellTiter-Glo Luminescent Cell Viability Assay (Promega) and crystal violet staining. For CellTiter-Glo assays, KP and KPK cells were seeded into 96-well white-walled plates at a density of 2,500 cells per well. The following day, cells were treated with vehicle (DMSO), IXA4 (10 µM), thapsigargin (Tg, 0.5 µM), BSO (50 µM) or the IRE1 RNase inhibitor 4µ8c (32 µM) as indicated. After 3 days, CellTiter-Glo was added to each well according to the manufacturer’s instructions. Luminescence was measured using a plate reader. Data are presented as viability normalized to vehicle-treated controls. For crystal violet staining, cells were seeded in 6-well plates and treated with compounds for 2 days. Media was removed, cells were washed with DPBS, and fixed with 100% methanol for 10 min. Fixed cells were stained with 0.5% crystal violet solution for 20 min at room temperature. Excess stain was washed away with water, and plates were dried and imaged.

### Electron microscopy preparation and data processing

#### Cloning, Expression and Purification of SLC33A1

Full length human SLC33A1(1-549, Uniprot ID: O00400 ACATN_HUMAN) was cloned into pCDNA3.1 by PCR with an N-terminal 6xHis-Flag-tag followed by a 3C cleavage sequence. Plasmid was amplified in DH5α cells and purified using Maxi prep kit (ZymoPURE II Plasmid Maxiprep Kit). 6×His-Flag-3C-SLC33A1 was expressed in Expi293-GnTI-cells. Transient transfection was performed in 1000 ml of medium (Expi293TM Expression Medium) in 3 L flasks using PEI reagent following the manufacturer’s protocol. 24 h prior to transfection, Expi293-GnTI-cells were split to a density of ∼2×10^6^ cells/mL, ensuring cells in log-phase growth and viability is above 95%. On the day of transfection, cells were adjusted to a density of ∼2.7-3.0×10^6^ cells/mL. For 1L of cells, 1 mg of the SLC33A1 DNA was diluted into a final volume of 25 mL Opti-MEM reduced-serum medium (Thermo Fisher Scientific) and filtered using a 0.22 µm filter assembly, similarly PEI at six times the amount of DNA was diluted separately in 25 mL of Opti-MEM reduced-serum medium and filtered. The sterile DNA and PEI mixtures were combined and incubated for 20 min at room temperature, before being added to the cell culture dropwise. Cells were incubated on an orbital shaking platform at 37°C with 8% CO_­_. After 20 h of transfection, sterile-filtered valproic acid (5 mM final concentration; Sigma) and sodium butyrate (5 mM final concentration) were added to the cells. From one liter expression, around 10 g cells were harvested 72 h after transfection by centrifugation (1500*g* for 20 min, 4°C) and stored at -80°C.

### Protein Purification

Cells were resuspended in Low Salt Buffer (20 mM Tris, pH 8.0, 150 mM NaCl) at a ratio of 5 mL per g of cell pellet. The suspension was homogenized using 30–50 strokes of a tight-fitting Dounce homogenizer. The lysate was centrifuged at 100,000*g* for 45 min at 4°C, and the resulting pellet was collected. The pellet was then resuspended in high salt buffer (20 mM Tris, pH 8.0, 600 mM NaCl) at a ratio of 5 mL per g of pellet and homogenized using 30-50 strokes of the Dounce homogenizer. The homogenate was centrifuged at 100,000*g* for 45 min at 4 °C, and the pellet was collected. The final pellet was resuspended in membrane protein extraction buffer (20 mM Tris, pH 8.0, 150 mM NaCl, 1.5% DDM, 0.1% CHS, and Roche One protease inhibitor cocktail tablet per 50 mL of buffer) at a ratio of 5 mL per g of pellet. The suspension was shaken overnight at 4°C to fully solubilize the membrane protein. The lysate was then centrifuged at 100,000*g* for 45 min at 4°C, and the resulting supernatant was collected and incubated with Ni-NTA resin pre-equilibrated with wash buffer (20 mM Tris, pH 8.0, 150 mM NaCl, 0.01%/0.001% LMNG/CHS). The resin was washed thoroughly, and the protein was eluted using elution buffer (20 mM Tris, pH 8.0, 150 mM NaCl, 0.01%/0.001% LMNG/CHS, 250 mM imidazole). The pooled fractions from the IMAC purification step were applied to anti-DYKDDDDK G1 affinity resin pre-equilibrated with Wash Buffer (20 mM Tris, pH 8.0, 150 mM NaCl, 0.01%/0.001% LMNG/CHS). The resin was washed, and the protein was eluted using Elution Buffer (20 mM Tris, pH 8.0, 150 mM NaCl, 0.01%/0.001% LMNG/CHS, 250 µg/mL Flag peptide). The protein from the Flag affinity chromatography (AC) step was concentrated and subjected to SEC using a Superdex 200 Increase 10/300 GL column (24 mL) equilibrated with SEC Buffer (20 mM Tris, pH 8.0, 150 mM NaCl, 0.005%/0.0005% LMNG/CHS). SEC peak fractions were pooled and concentrated to 2.2 mg/mL (A_­_) using Millipore Amicon Ultra Centrifugal Filters (15 mL, 30 kDa cutoff).

### Nanodisc reconstitution

For Liposome Preparation, lipids (POPC:DOPE:PS = 55:30:15 molar ratio) were dissolved in chloroform and dried under nitrogen gas for 2 h, then further dried under vacuum for 4 h. The lipid film was rehydrated in 20 mM Tris, pH 8.0, 150 mM NaCl to obtain a 10 mM liposome suspension. This was clarified using freeze-thaw cycles followed by bath sonication until a milky suspension was achieved. To destabilize the liposomes, 0.047% LMNG/CHS was added and incubated at room temperature for 3 h. Purified SLC33A1 was mixed with the destabilized liposomes at 4 °C for 30 min, followed by the addition of MSP1D1 purified in house following at 4°C for 10 min^52^. AMBERLIFETM XAD-2 beads (0.8 g wet beads/mL) were then added and rotated overnight at 4°C to remove free detergents. The protein: MSP1D1: liposome ratio was maintained at a molar ratio of 1:5:250. The reconstituted nanodiscs were subjected to Flag AC using 5 mL of Flag resin pre-equilibrated with Wash Buffer (20 mM Tris, pH 8.0, 150 mM NaCl). The resin was washed and eluted with Elution Buffer (20 mM Tris, pH 8.0, 150 mM NaCl, 250 µg/mL Flag peptide).

The nanodiscs from the Flag AC step were subjected to SEC using a Superdex 200 Increase 10/300 GL column (24 mL) equilibrated with SEC Buffer (20 mM Tris, pH 8.0, 150 mM NaCl). SEC fractions were pooled and concentrated to 1.85 mg/mL (A_­_) using Millipore Amicon Ultra Centrifugal Filters (15 mL, 30 kDa cutoff).

### Cryo-EM sample preparation and data collection

3 µLof concentrated sample (5 mg/mL) was applied to freshly glow-discharged R0.6/1 UltrAuFoil grids (Quantifoil). For drug bound 100 µM of IXA4 was incubated with protein for 10 min on ice. Grids were blotted for 5 s and plunge frozen in liquid ethane using a manual plunger in a 4°C cold room with >95% humidity. Movies were collected using a Talos Arctica transmission electron microscope (Thermo Fisher Scientific) operating at 200 keV under parallel illumination conditions. Micrographs were acquired with a Falcon 4i direct electron detector (Thermo Fisher Scientific) at a nominal magnification of 190,000×, corresponding to a calibrated pixel size of 0.727 Å/pixel at the specimen level with a total accumulated dose of ∼40 e⁻/Å² and the data was saved in the electron-event representation (EER) format. Automated data acquisition was performed using EPU software (version 3.9.1.8206; Thermo Fisher Scientific) with an aberration-free image shift (AFIS) strategy and a maximum image shift of up to 8 µm. Defocus values were systematically varied from -0.8 to -2.4 µm in increments of 0.4 µm. Detailed dataset parameters, including total micrographs (for Apo and IXA4-bound SLC33A1), are provided in **Table S3**.

### Cryo-EM data processing and image analysis

The processing workflow used for image analysis of both datasets is shown in supplementary figure (**Fig. S3**, **S4**). For apo SLC33A1 all image processing was performed using cryoSPARC (version v4.7.0)^53^. Accepted exposures with defocus range from 0.8-2.4 µm and contrast transfer function (CTF) cut off < 5 Å were motion corrected (patch based) and CTF was estimated in cryo-SPARC live. After each session, the accepted exposures and particles from the cryoSPARC Live session were exported into the respective cryoSPARC workspace. For apo SLC33A1 6,925 micrographs after pre-processing were used for particle picking using blob picker to select particles with diameters ranging from 80 to 110 Å and later a template was generated from live 2Ds which was used to re-pick particles using template picker. Around 4 million particles were extracted with a box size of 256 pixels Fourier-crop binned to 100 pixels. Iterative rounds of 2D classification were done for an initial clean up resulting in 1.2 million particles. Selected 2D classes were used to generate two ab-initio volumes with 0 class similarity, initial resolution 12 Å, maximum resolution 8 Å, number of initial iterations 300, number of final iterations 400, initial mini batch size 300, final mini batch size 1200. These ab-initio were used as input for the heterogeneous refinement by using all the particles from selected 2Ds and classifying them on these ab-initio keeping all default parameters for heterogeneous refinement job with C1 symmetry. Cleaned particles (383,455) from this hetero refinement were used to generate 2X parallel ab-initio jobs, with 0 class similarity, two classes each, initial low pass resolution 9 Å, maximum resolution 6 Å. Particles from good classes of hetero refinement were extracted and re-centered using alignment shifts with full box size of 256 pixels. Non-uniform refinement was performed with low pass resolution 6, keeping minimize over per particle scale and initial noise model from images true, number of extra final poses 1, per-particle defocus, and per group CTF parameters true. After global and local CTF refinement, an initial mask encompassing the transmembrane helices and soluble domain was generated in UCSF ChimeraX by converting an AlphaFold Server model to a low-resolution density using the molmap command (15 Å)^54^. The AlphaFold model was obtained from AlphaFold Server^55^. The resulting volume was imported into cryoSPARC using “Import 3D volume” job and refined using the “Volume Tools” job (threshold 0.045, dilation radius 2 pixels, soft padding 15 pixels) to generate the final mask used for local refinement. Local refinement was performed with recenter rotation and shift each iteration enabled, using a rotation search extent of 6° and a shift search extent of 3 Å, with an initial low-pass resolution of 5 Å resulting to 3.3 Å final map. The refined map was post-processed with EMready2 and local resolution was estimated in cryoSPARC and is shown in Fig. S3^56^. The image-processing workflow for the IXA4-bound SLC33A1 dataset is summarized in **Fig. S4**. CTF estimation (patch CTF) and motion correction was performed using live cryosparc with default parameters, and 6,486 micrographs with estimated CTF resolutions better than 5 Å were retained. Particles were initially picked using a blob picker, and representative classes from an initial round of live 2D classification were used as templates for template-based particle picking with a particle diameter range of 80-110 Å. Using template picking, 3,119,666 particles were extracted with a 256-pixel box size and 2.5× Fourier binning. Multiple rounds of 2D classification were performed to remove junk and poorly aligned particles. Selected 2D classes were used to generate two ab initio volumes with class similarity set to zero, using an initial resolution of 9 Å and a maximum resolution of 6 Å. These volumes were generated with 300 initial and 400 final iterations, with the mini-batch size increased from 300 initial to 1200 final. The resulting ab initio models were used as references for heterogeneous refinement into three classes, separating one well-resolved class from two poorly resolved classes. A total of 457,520 particles from the well-resolved class were selected, re-extracted at the full box size, and refined using non-uniform refinement with an initial low-pass filter of 6 Å, enabling per-particle defocus refinement, per-group CTF refinement, optimization of per-particle scale, and refinement of one additional final pose. This refinement yielded a reconstruction at 3.4 Å resolution. Following global and local CTF refinement, a soft mask encompassing the transmembrane helices was generated in UCSF ChimeraX by converting the apo model into a low-resolution density using the molmap command at 15 Å. The resulting volume was imported into cryoSPARC and processed using the Volume Tools job to generate the final refinement mask. Local refinement was then performed with recentering of rotation and translation at each iteration, resulting in a final reconstruction at 3.27 Å resolution. The map was post-processed using EMReady2, and local resolution estimation was performed in cryoSPARC (**Fig. S4**).

### Atomic model building and refinement

Model building and refinement were initiated with AlphaFold model^55^. Restraints and PDB for IXA4 were generated with Phenix tool eLBOW^57^. Iterative rounds of model building and refinement were performed in PHENIX v2.0orc1-5617 and Coot 0.9.8.95 EL until a reasonable agreement between the model and data was achieved^58,59^. Phenix real space refinement includes global minimization, rigid body, local grid search, atomic displacement parameters, and morphing for the first cycle. It was run for 100 iterations, five macro cycles, with a target bonds RMSD of 0.01 Å and a target angles RMSD of 1.0°. The refinement settings also include the secondary structure restraints and Ramachandran restraints. ChimeraX plug-in ISOLDE^60^ was used to manually fix any Ramachandran outliers, rotamers, and clashes not corrected by Phenix. The map and model were validated with Molprobity^61^ within Phenix. In some cases, maps modified with the EMready2 tool were used to aid in modeling. ChimeraX was used to interpret the EM reconstructions and atomic models, as well as to generate figures. PoseView^62^ was utilized to analyze the molecular interactions between the ligand and the protein binding site, while MOLEonline^36^ was employed to visualize the potential translocation pathway of the ligand through the transporter.

### LC/MS analysis for whole-cell and ER-isolated polar metabolites

The ER was purified from HEK293T cells expressing EMC3-mScarlet-3xHA (ER-tag) or EMC3-mScarlet-3xMYC (control-tag), as previously described^28^. Briefly, cells cultured in two 15 cm dishes were treated with DMSO or IXA4 (10 µM) for 4 h. Cells were then harvested, rinsed twice with chilled saline (0.9% NaCl), and scraped into 3 mL of cold KPBS before being pelleted at 1,500*g* for 2 min at 4°C. After resuspending the pellet in 1 mL of KPBS, two 10 µL aliquots were reserved: one was lysed in 90 µL of 1% (v/v) Triton lysis buffer (supplemented with protease inhibitors) for whole-cell protein analysis, while the other was transferred into 90 μµl of an ACN:MeOH:H2O (2:2:1) mixture containing heavy-labeled amino acid standards for direct whole-cell metabolite extraction. The remaining cell suspension was homogenized using a 2 mL Dounce homogenizer via two sets of 30 strokes. To isolate the ER, the resulting homogenate was cleared by centrifugation at 3,000*g* for 10 min at 4°C, and the supernatant was subsequently incubated with pre-washed anti-HA magnetic beads for 5 min at 4°C. Following three washes with cold KPBS, 10% of the beads were processed for protein quantification, while the remaining 90% underwent metabolite extraction in 50 µL of ACN:MeOH:H2O (2:2:1) containing heavy labeled amino acid standards via 10 s of vortexing followed by 10 min of rotation at 4°C. After removing the beads with a magnet and performing a final clearing spin at 20,000*g* for 15 min, the samples were analyzed via LC-MS polar metabolite profiling without drying. Metabolomic data were normalized to total protein (BCA) or NAD^+^ abundance for whole-cell samples, and to UDP-GlcNAc/GalNAc levels for ER-specific extracts. Polar metabolite profiling was performed using a Thermo Scientific Vanquish Flex UHPLC system coupled to an Orbitrap ID-X mass spectrometer. Chromatographic separation was achieved at 45°C on a HILICON iHILIC-(P) Classic column (100 x 2.1 mm, 5 µm) protected by a matching guard column (20 x 2.1 mm, 5 µm). The mobile phase was composed of 20 mM ammonium bicarbonate, 2.5 µM medronic acid, and 0.1% ammonium hydroxide in 95:5 H2O:ACN (Solvent A) and 95:5 ACN:H2O (Solvent B). A 4 µL sample was injected and eluted at a flow rate of 250 µL/min using the following linear gradient: 0-1 min at 90% B; 12 min at 35% B; 12.5-14.5 min at 25% B; and returning to 90% B at 15 min. Mass spectra were collected in both positive and negative ionization modes. The electrospray ionization source was operated with a spray voltage of 3.5 kV (positive) and -2.8 kV (negative), while gas settings were maintained at 50 (sheath), 10 (auxiliary), and 1 (sweep). The temperatures for the ion transfer tube and vaporizer were set to 300°C and 200°C, respectively. Full scan and MS/MS data were acquired across a mass range of 67–1000 Da with resolutions of 120,000 (MS1) and 30,000 (MS/MS). Maximum injection times were 200 ms for MS1 and 100 ms for MS/MS, utilizing a 1.5 Da isolation window. Finally, raw LC/MS data were processed and integrated using a combination of XCMS, Compound Discoverer, and Skyline.

## ACKNOWLEDGEMENTS

This work was funded by the National Institutes of Health (AG046495 to RLW, DK137470 to ES and RLW, UM1 TR004407 to ES and RLW, GM069832 to SF). SK was supported by a PhRMA Foundation predoctoral fellowship. MR and PB were supported by Hewitt Foundation postdoctoral fellowships. SL was supported by the Human Frontier Science Program. We thank Richard Labaudiniere, Steven Wilkens, Bo Qin, and Mack Flinspach for their input and feedback.

---

**Figure S1.**
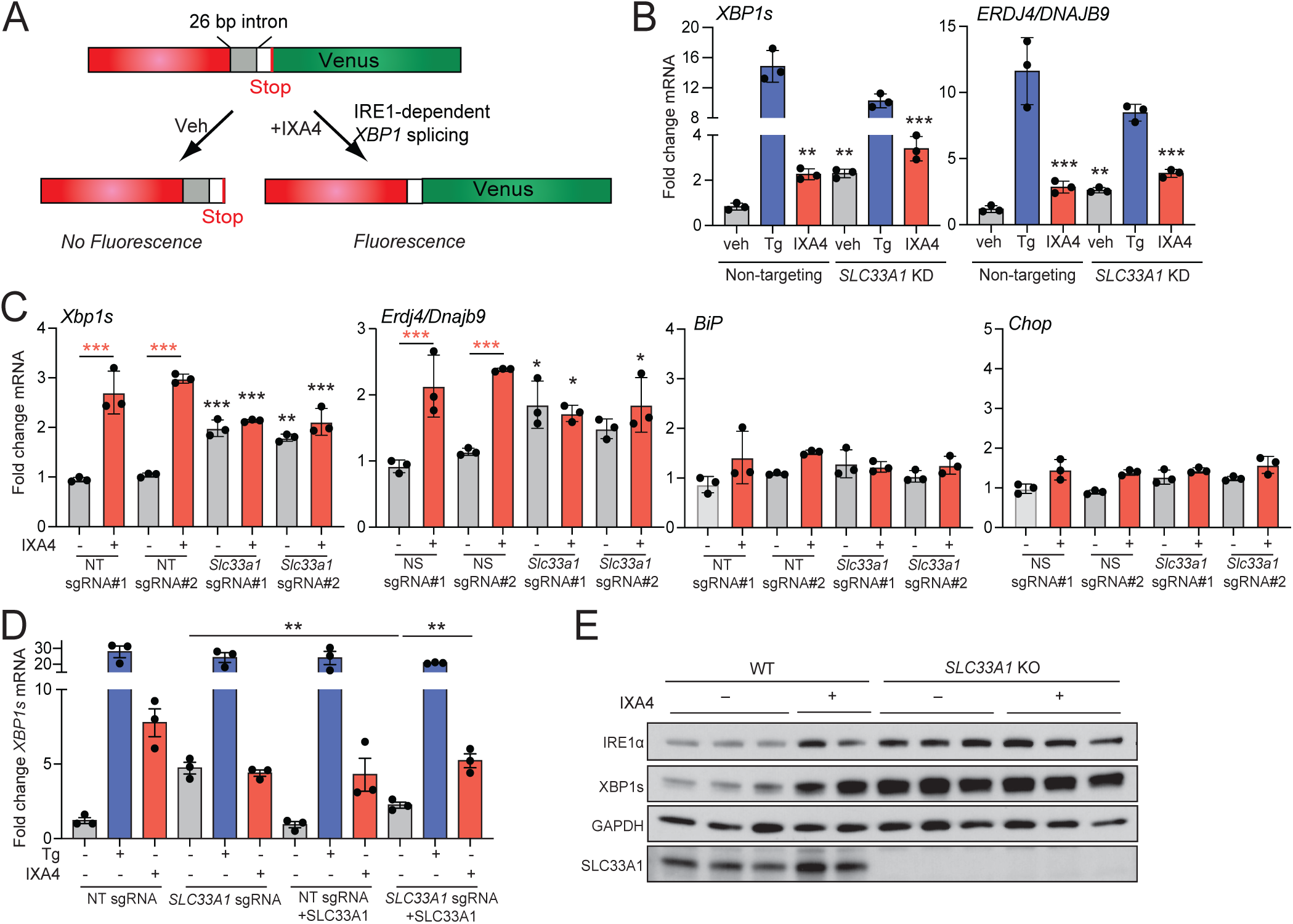
(Related to Figure 1). CRISPR screen identifies SLC33A1 as a protein involved in IXA4-induced IRE1/XBP1s signaling. **A.** Schematic of the XBP1-Venus IRE1 splicing reporter. **B**. Expression, measured by qPCR, of the IRE1/XBP1s target genes *XBP1s* and *ERDJ4/DNAJB9* in HEK293 cells expressing non-targeting or *SLC33A1* shRNA and treated for 4 h with Tg (0.5 µM) or IXA4 (10 µM). **C**. Expression, measured by qPCR, of the IRE1 target genes *Xbp1s* and *Erdj4/Dnajb9*, the ATF6 target gene *BiP*, and the PERK target gene *Chop* in MEF cells CRISPR-deleted of *Slc33a1* using two distinct sgRNA and treated for 4 h with IXA4 (10 µM). MEF cells expressing two distinct non-targeting sgRNAs are shown as a control. **D**. Expression, measured by qPCR, of the IRE1 target gene *XBP1s* in non-targeting HEK293 cells or HEK293 cells lacking *SLC33A1* transfected with mock or wild-type SLC33A1, as indicated, and then treated for 4 h with Tg (0.5 µM) or IXA4 (10 µM). **E**. Immunoblots of the indicated proteins in HEK293T cells expressing non-targeting or *SLC33A1* sgRNAs treated for 4 h with IXA4 (10 µM). Black asterisks in (**C**) represent comparisons between vehicle-treated control cells expressing NT sgRNA #1 and mutant cells treated with or without IXA4. Red asterisks in (**C**) indicate comparisons between vehicle and IXA4-treated samples. *p<0.05, **p<0.01, ***p<0.005, one-way ANOVA.

**Figure S2.**
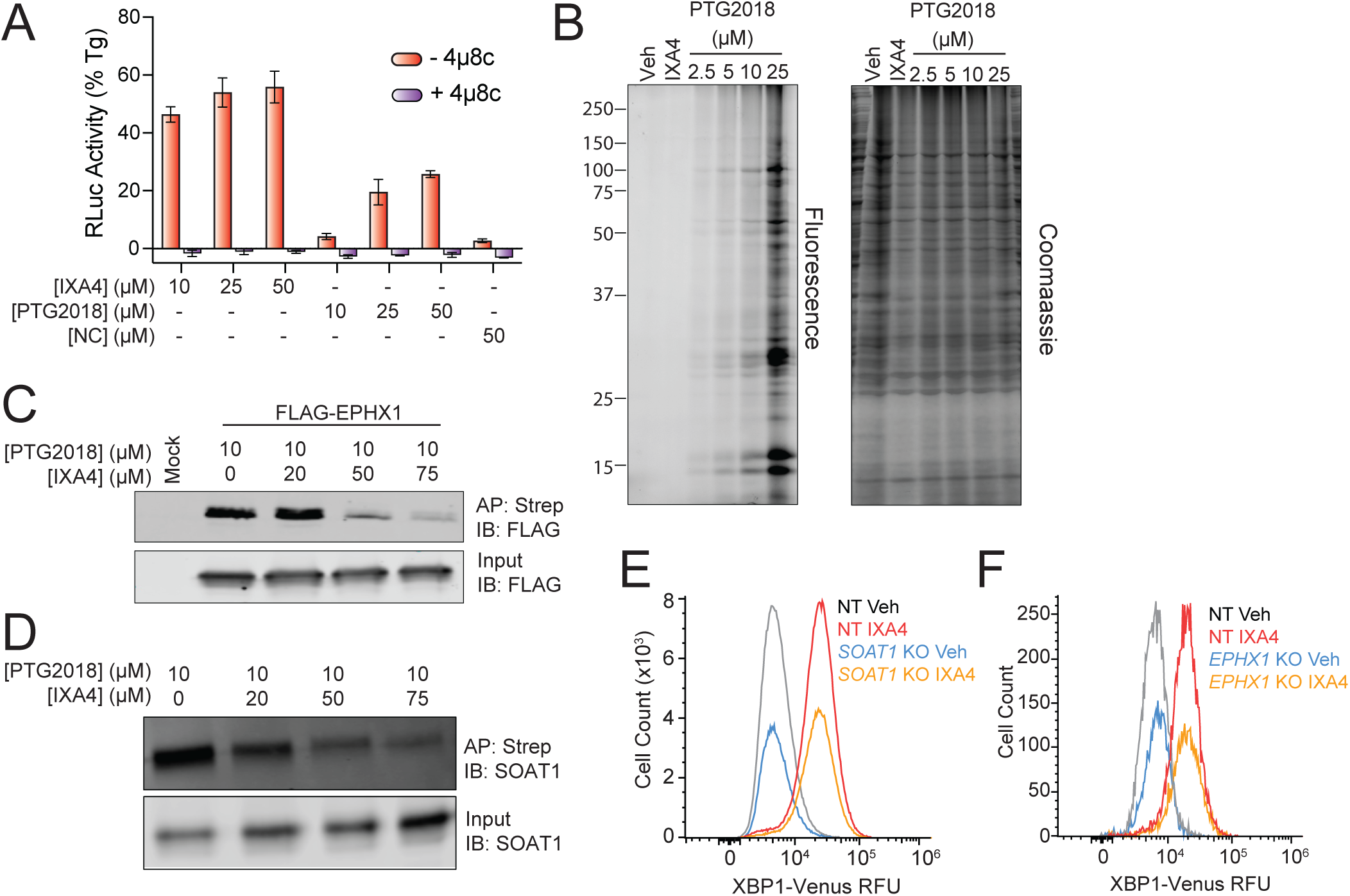
(Related to Figure 2). IXA4 binds SLC33A1. **A.** XBP1-Rluc signal in HEK293 cells stably expressing XBP1-RLuc treated with the indicated concentration of IXA4 or PTG2018 in the presence or absence of the IRE1 RNAse inhibitor 4µ8c (32 µM) for 14 h. Data are normalized to the XBP1-RLuc signal seen in cells treated with thapsigargin (Tg; 0.5 µM). Error bars show SEM for n=3 replicates. **B**. Dose-dependent proteome labeling with PTG2018 (left, rhodamine fluorescence; right, Coomassie stain). PTG2018-labeled proteins were identified by appending TAMRA-azide to PTG2018 via click chemistry. **C**. Streptavidin affinity purification and proteomes (input) from HEK293T cells overexpressing FLAG-tagged EPHX1 and treated with the indicated concentration of PTG2018 and IXA4. PTG2018 labeling of FLAG-tagged EPHX1 was visualized by appending a biotin to the alkyne handle of PTG2018 using click chemistry and monitoring the recovery of FLAG-tagged protein in streptavidin isolates. Mock transfected cells are shown as a control. **D**. Streptavidin affinity purification and proteomes (input) from HEK293T cells treated with the indicated concentration of PTG2018 and IXA4. PTG2018 labeling of SOAT1 was visualized by appending biotin to the alkyne handle of PTG2018 via click chemistry and monitoring the recovery the recovery of endogenous SOAT1 in streptavidin isolates. **E**, **F**. XBP1-Venus signal, measured by flow cytometry, in HEK293 cell stably expressing Cas9 and transduced with lentivirus encoding non-targeting (NT), *SOAT1 or EPHX1* sgRNA treated for 14 h with vehicle or IXA4 (10 µM).

**Figure S3.**
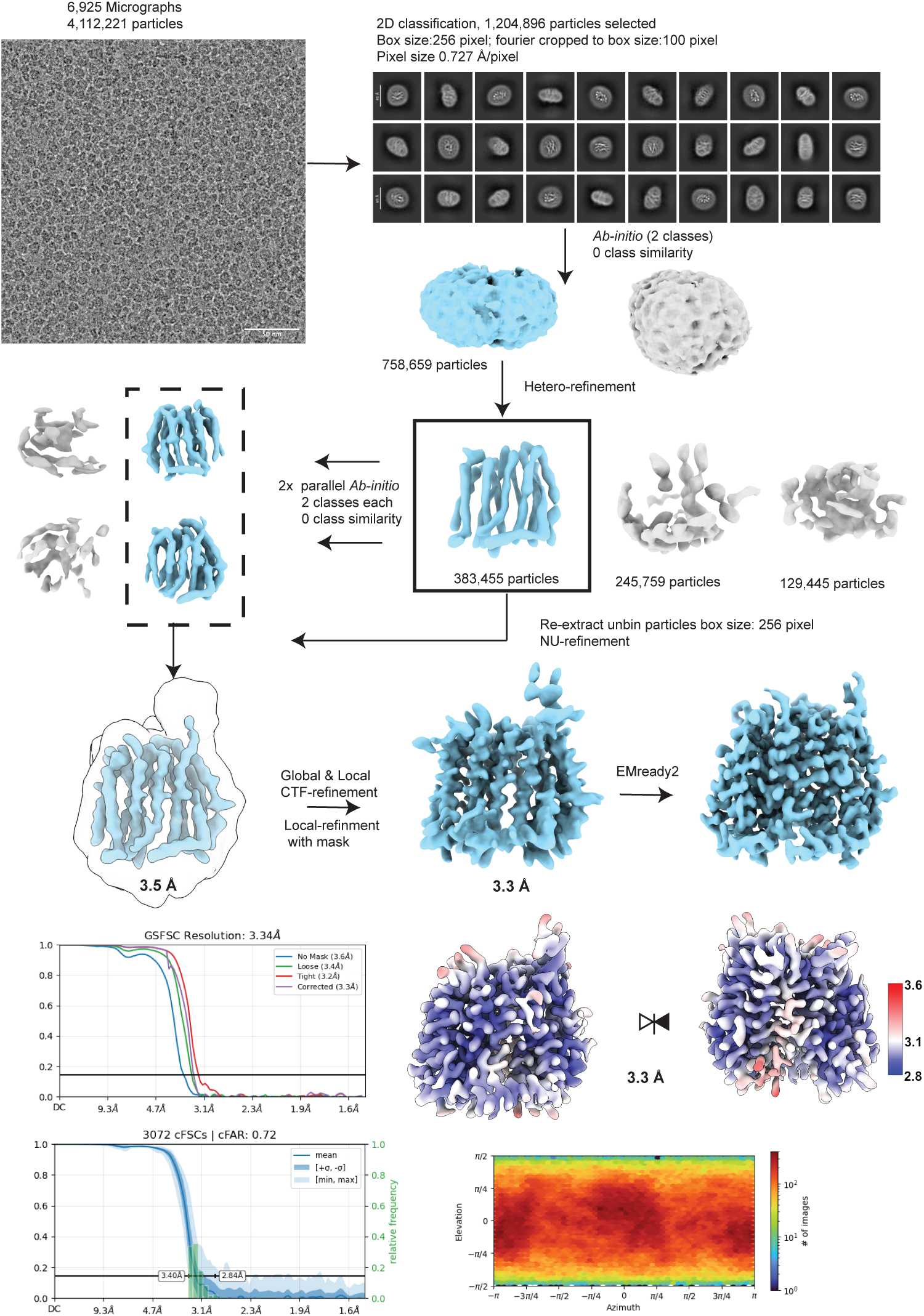
(Related to Figure 3). Cryo-EM image-processing workflows for apo SLC33A1. Image-processing workflow for apo SLC33A1. Representative micrographs with 5Å applied lowpass and 2D class averages are shown. Selected particles from good classes were used to generate ab initio reconstructions (two classes) and subsequently refined by heterogeneous refinement to isolate a well-resolved class. In parallel, two additional ab initio reconstructions (two classes each) were performed, and the best-resolved volume was used as the reference for downstream refinement. Particles from the well-resolved heterogeneous-refinement class were then re-extracted to full box size and refined by non-uniform refinement, followed by global and local CTF refinement and masked local refinement. The final reconstruction was post-processed with EMReady2, and validation outputs including gold-standard FSC, cFSC, angular distribution, and local-resolution estimation in Å are shown.

**Figure S4.**
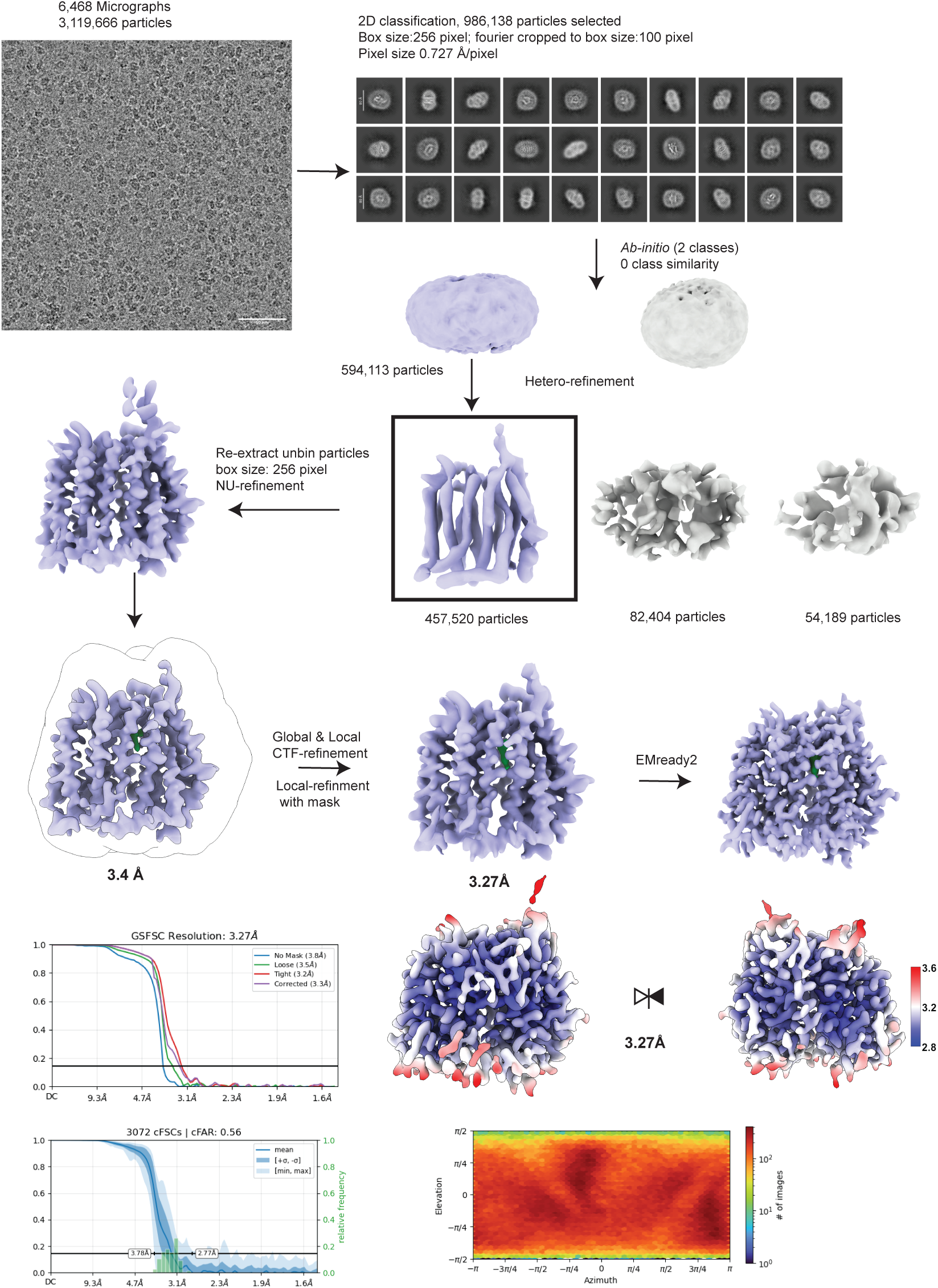
(Related to Figure 3). Cryo-EM image-processing workflows for IXA4-bound SLC33A1. Image-processing workflow for IXA4-bound SLC33A1. Representative micrographs with 5Å applied lowpass and 2D class averages are shown. Particles from selected 2D class averages were used to generate ab initio models, which served as references for heterogeneous refinement. Particles from the well-resolved class were re-extracted and refined by non-uniform refinement, followed by global and local CTF refinement and masked local refinement. The final map was post-processed with EMReady2. Corresponding gold-standard FSC, cFSC, angular distribution, and local-resolution estimation in Å are shown.

**Figure S5.**
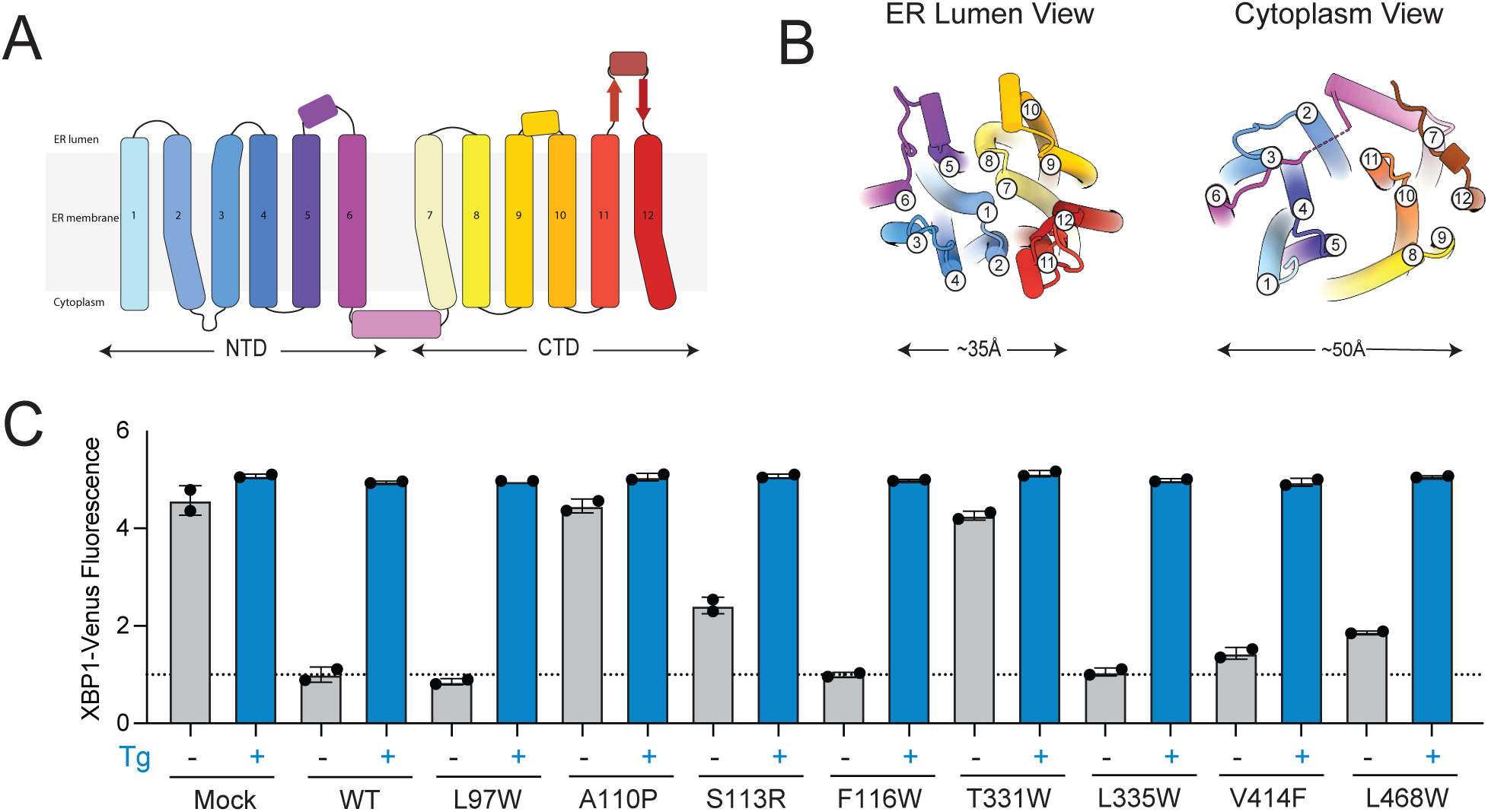
(Related to Figure 3). IXA4 binds the central channel of SLC33A1. **A**. Topology diagram of SLC33A1 showing 12 transmembrane helices (TM1-TM12) and their orientation in the ER membrane. TM helices are colored from N to C terminus as light blue, cornflower blue, dodger blue, steel blue, violet, magenta, lemon chiffon, yellow, gold, orange, tomato, and firebrick. The lateral helix is colored plum, and the terminal segment is colored saddle brown. **B.** Ribbon representations of SLC33A1 in the ER-lumen view (left) and the cytoplasmic view (right), illustrating the closed and open conformations, respectively. Transmembrane helices are colored and numbered as in panel **A**. **C.** XBP1-Venus signal, measured by flow cytometry, after reconstitution of wild-type SLC33A1 and mutant proteins in HEK293 XBP1-Venus reporter cells lacking endogenous SLC33A1 following treatment with vehicle or Tg (0.5 µM) for 14 h.

**Figure S6.**
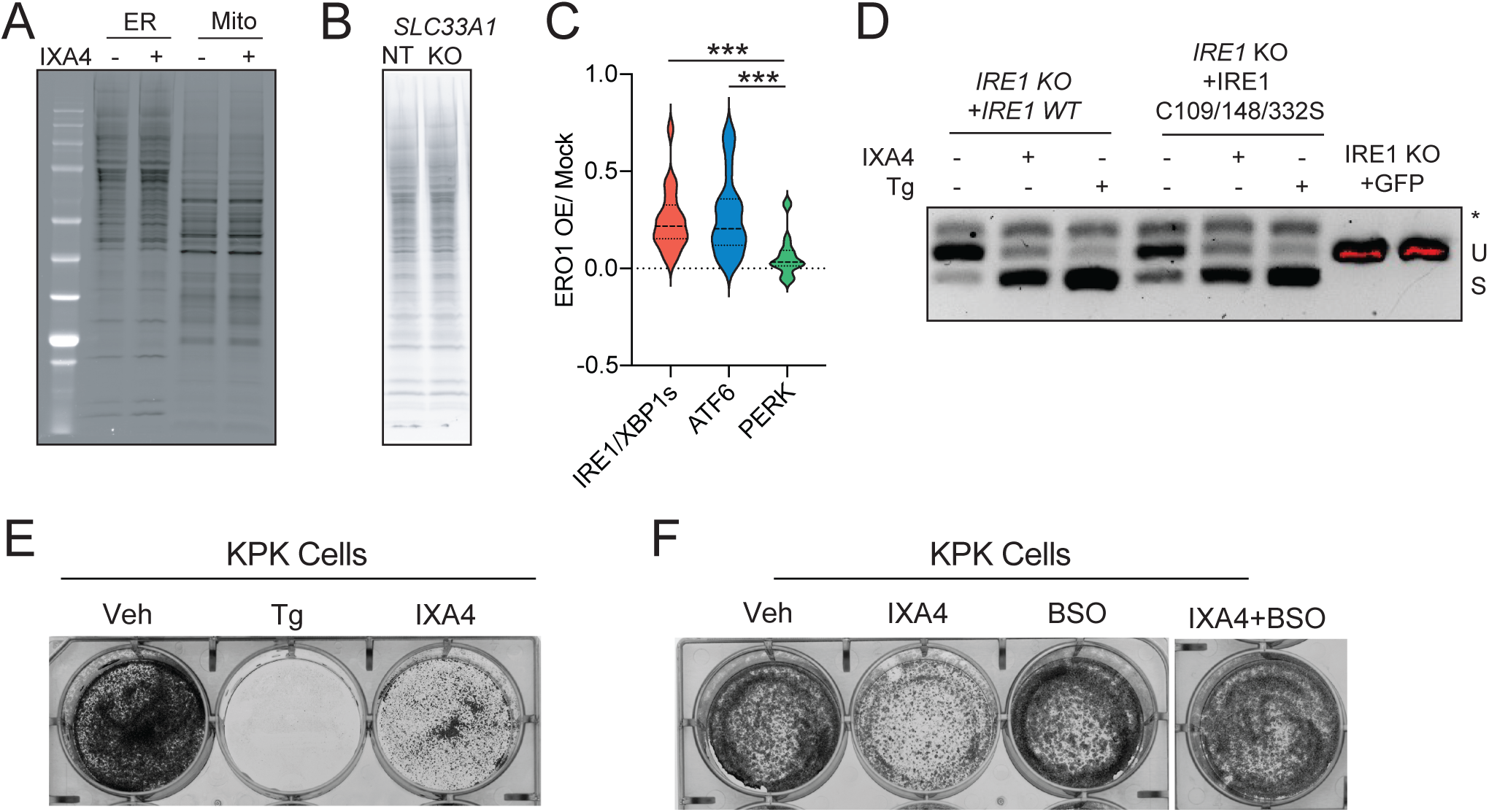
(Related to Figure 4). IXA4 binding to SLC33A1 promotes ER hyperoxidation and IRE1 activation. **A**. In-gel fluorescence of acetylated proteins in ER and mitochondrial fractions isolated from HEK293T cells treated for 3 h with IXA4 (10 µM) and then incubated for 1 h with 3-butynoic acid. Acetylated proteins were identified by appending a rhodamine tag onto the 3-butynoic acid probe via click chemistry. **B**. In-gel fluorescence of acetylated proteins in ER fractions isolated from HEK293T Cas9 cells expressing non-targeting (NT) or *SLC33A1* sgRNA incubated for 1 h with the 3-butynoic acid probe. Acetylated proteins were then identified by appending a rhodamine tag onto the 3-butynoic acid probe via click chemistry. **C**. Expression (published DNA microarrays)^38^ of IRE1/XBP1s, ATF6, and PERK target genesets^48^ in HEK293T cells overexpressing the ER oxidase ERO1. Source data available in GEO (GSE40601). **D**. *XBP1* mRNA splicing (RT-PCR) in *IRE1*-deficient HeLa cells transfected with wild-type IRE1 or a C109S/C148S/C332S IRE1 triple mutant and treated for 2 h with IXA4 (10 µM) or thapsigargin (Tg; 0.5 µM). Unspliced (u), spliced (s), and hybrid (*), *XBP1* are shown. **E**. Crystal violet stained KPK cells treated for 2 days with thapsigargin (Tg; 0.5 µM) or IXA4 (10 µM). **F**. Crystal violet stained KPK cells incubated for 2 days with IXA4 (10 µM) and/or BSO (50 µM). p<0.05, **p<0.01, ***p<0.005, one-way ANOVA.

## SUPPLEMENTAL TABLE LEGENDS

**Table S1 (related to Figure 1). CRISPR screen identifies SLC33A1 as a protein involved in IXA4-induced IRE1/XBP1s signaling**

Phenotypes, fold change and pvalue (Table S1A) and counts (Table S1B) for genes targeted in the CRISPR screen for DMSO and IXA4-treated cells were analyzed using MAGeCK (see Methods for details).

**Table S2 (related to Figure 2). IXA4 binds SLC33A1**

Fold change and pvalue with a corresponding competition excess (Table S2A) and TMT data (Table S2B) for proteins identified a competition chemoproteomics experiment for IXA4 and PTG2018-treated cells

**Table S3 (related to Figure 3). IXA4 binds the central channel of SLC33A1**

Cryo-EM data collection, refinement, and validation

**Table S4 (additional materials).** Table showing the primers used to for qPCR and RT-qPCR, and protospacers used for CRISPR in this manuscript.

## REFERENCES

1. Farrugia, M.A. & Puglielli, L. Nepsilon-lysine acetylation in the endoplasmic reticulum - a novel cellular mechanism that regulates proteostasis and autophagy. J Cell Sci 131(2018).

2. Hirabayashi, Y., Kanamori, A., Nomura, K.H. & Nomura, K. The acetyl-CoA transporter family SLC33. Pflugers Arch 447, 760–2 (2004).

3. Jonas, M.C., Pehar, M. & Puglielli, L. AT-1 is the ER membrane acetyl-CoA transporter and is essential for cell viability. J Cell Sci 123, 3378–88 (2010).

4. Chiplunkar, S. et al. Huppke-Brendel syndrome in a seven months old boy with a novel 2-bp deletion in SLC33A1. Metab Brain Dis 31, 1195–8 (2016).

5. Parayil Sankaran, B. et al. Huppke-Brendel Syndrome. in GeneReviews((R)) (eds. Adam, M.P. et al.) (Seattle (WA), 1993).

6. Lin, P. et al. A missense mutation in SLC33A1, which encodes the acetyl-CoA transporter, causes autosomal-dominant spastic paraplegia (SPG42). Am J Hum Genet 83, 752–9 (2008).

7. Mao, F. et al. Identification and functional analysis of a SLC33A1: c.339T>G (p.Ser113Arg) variant in the original SPG42 family. Hum Mutat 36, 240–9 (2015).

8. Luo, Y. et al. Mutation and clinical characteristics of autosomal-dominant hereditary spastic paraplegias in China. Neurodegener Dis 14, 176–83 (2014).

9. Lin, P. et al. Prenatal diagnosis of autosomal dominant hereditary spastic paraplegia (SPG42) caused by SLC33A1 mutation in a Chinese kindred. Prenat Diagn 30, 485–6 (2010).

10. Hullinger, R. et al. Increased expression of AT-1/SLC33A1 causes an autistic-like phenotype in mice by affecting dendritic branching and spine formation. J Exp Med 213, 1267–84 (2016).

11. Romero, R. et al. Keap1 mutation renders lung adenocarcinomas dependent on Slc33a1. Nat Cancer 1, 589–602 (2020).

12. Peng, Y. et al. Improved proteostasis in the secretory pathway rescues Alzheimer’s disease in the mouse. Brain 139, 937–52 (2016).

13. Grandjean, J.M.D. et al. Pharmacologic IRE1/XBP1s activation confers targeted ER proteostasis reprogramming. Nat Chem Biol 16, 1052–1061 (2020).

14. Hetz, C., Zhang, K. & Kaufman, R.J. Mechanisms, regulation and functions of the unfolded protein response. Nat Rev Mol Cell Biol 21, 421–438 (2020).

15. Acosta-Alvear, D., Harnoss, J.M., Walter, P. & Ashkenazi, A. Homeostasis control in health and disease by the unfolded protein response. Nat Rev Mol Cell Biol 26, 193–212 (2025).

16. Hetz, C. The unfolded protein response: controlling cell fate decisions under ER stress and beyond. Nat Rev Mol Cell Biol 13, 89–102 (2012).

17. Grandjean, J.M.D. & Wiseman, R.L. Small molecule strategies to harness the unfolded protein response: where do we go from here? J Biol Chem 295, 15692–15711 (2020).

18. Marciniak, S.J., Chambers, J.E. & Ron, D. Pharmacological targeting of endoplasmic reticulum stress in disease. Nat Rev Drug Discov 21, 115–140 (2022).

19. Madhavan, A. et al. Pharmacologic IRE1/XBP1s activation promotes systemic adaptive remodeling in obesity. Nat Commun 13, 608 (2022).

20. Touvier, T. et al. Activation of XBP1s attenuates disease severity in models of proteotoxic Charcot-Marie-Tooth type 1B. Brain 148, 1978–1993 (2025).

21. Li, Y., Liu, D. & Li, S. IRE1/Xbp1 promotes the clearance of poly(GR) dipeptide repeats in amyotrophic lateral sclerosis. J Biol Chem 301, 110764 (2025).

22. Dieterich, I.A. et al. Acetyl-CoA flux regulates the proteome and acetyl-proteome to maintain intracellular metabolic crosstalk. Nat Commun 10, 3929 (2019).

23. Iwawaki, T., Akai, R., Kohno, K. & Miura, M. A transgenic mouse model for monitoring endoplasmic reticulum stress. Nat Med 10, 98–102 (2004).

24. Doench, J.G. et al. Optimized sgRNA design to maximize activity and minimize off-target effects of CRISPR-Cas9. Nat Biotechnol 34, 184–191 (2016).

25. Li, W. et al. MAGeCK enables robust identification of essential genes from genome-scale CRISPR/Cas9 knockout screens. Genome Biol 15, 554 (2014).

26. Tian, R. et al. CRISPR Interference-Based Platform for Multimodal Genetic Screens in Human iPSC-Derived Neurons. Neuron 104, 239–255 e12 (2019).

27. Ginto George, H.P.H., Richard Kay, David Ron, Adriana Ordóñez. Metabolite import via SLC33A1 enables ATF6 activation by endoplasmic reticulum stress. bioRxiv (2025).

28. Liu#, S., et al. SLC33A1 exports oxidized glutathione to maintain endoplasmic reticulum redox homeostasis. bioRxiv (2026).

29. Rostovtsev, V.V., Green, L.G., Fokin, V.V. & Sharpless, K.B. A stepwise huisgen cycloaddition process: copper(I)-catalyzed regioselective “ligation” of azides and terminal alkynes. Angew Chem Int Ed Engl 41, 2596–9 (2002).

30. Tornoe, C.W., Christensen, C. & Meldal, M. Peptidotriazoles on solid phase: [1,2,3]-triazoles by regiospecific copper(i)-catalyzed 1,3-dipolar cycloadditions of terminal alkynes to azides. J Org Chem 67, 3057–64 (2002).

31. Conway, L.P., Li, W. & Parker, C.G. Chemoproteomic-enabled phenotypic screening. Cell Chem Biol 28, 371–393 (2021).

32. Homan, R.A. et al. Photoaffinity labelling with small molecules. Nat Rev Methods Primers 30(2024).

33. Cross, B.C. et al. The molecular basis for selective inhibition of unconventional mRNA splicing by an IRE1-binding small molecule. Proc Natl Acad Sci U S A 109, E869–78 (2012).

34. Chiu, T.Y. et al. Chemoproteomic development of SLC15A4 inhibitors with anti-inflammatory activity. Nat Chem Biol 20, 1000–1011 (2024).

35. Zhou, D., Chen, N., Huang, S., Song, C. & Zhang, Z. Mechanistic insights into the acetyl-CoA recognition by SLC33A1. Cell Discov 11, 36 (2025).

36. Pravda, L. et al. MOLEonline: a web-based tool for analyzing channels, tunnels and pores (2018 update). Nucleic Acids Res 46, W368–W373 (2018).

37. Yang, Y.Y., Ascano, J.M. & Hang, H.C. Bioorthogonal chemical reporters for monitoring protein acetylation. J Am Chem Soc 132, 3640–1 (2010).

38. Hansen, H.G. et al. Hyperactivity of the Ero1alpha oxidase elicits endoplasmic reticulum stress but no broad antioxidant response. J Biol Chem 287, 39513–23 (2012).

39. Eletto, D., Chevet, E., Argon, Y. & Appenzeller-Herzog, C. Redox controls UPR to control redox. J Cell Sci 127, 3649–58 (2014).

40. Zavitsanou, A.M. et al. KEAP1 mutation in lung adenocarcinoma promotes immune evasion and immunotherapy resistance. Cell Rep 42, 113295 (2023).

41. Appenzeller-Herzog, C. Glutathione- and non-glutathione-based oxidant control in the endoplasmic reticulum. J Cell Sci 124, 847–55 (2011).

42. Ferdigg, A., Hopp, A.K., Wolf, G. & Superti-Furga, G. Membrane transporters modulating the toxicity of arsenic, cadmium, and mercury in human cells. Life Sci Alliance 8(2025).

43. Chakravarthi, S., Jessop, C.E. & Bulleid, N.J. The role of glutathione in disulphide bond formation and endoplasmic-reticulum-generated oxidative stress. EMBO Rep 7, 271–5 (2006).

44. Ushioda, R. & Nagata, K. Redox-Mediated Regulatory Mechanisms of Endoplasmic Reticulum Homeostasis. Cold Spring Harb Perspect Biol 11(2019).

45. Wang, L. & Wang, C.C. Oxidative protein folding fidelity and redoxtasis in the endoplasmic reticulum. Trends Biochem Sci 48, 40–52 (2023).

46. Cooley, M.M. et al. Deficient Endoplasmic Reticulum Acetyl-CoA Import in Pancreatic Acinar Cells Leads to Chronic Pancreatitis. Cell Mol Gastroenterol Hepatol 11, 725–738 (2021).

47. Shoulders, M.D. et al. Stress-independent activation of XBP1s and/or ATF6 reveals three functionally diverse ER proteostasis environments. Cell Rep 3, 1279–92 (2013).

48. Grandjean, J.M.D. et al. Deconvoluting Stress-Responsive Proteostasis Signaling Pathways for Pharmacologic Activation Using Targeted RNA Sequencing. ACS Chem Biol 14, 784–795 (2019).

49. Yau, E.H. & Rana, T.M. Next-Generation Sequencing of Genome-Wide CRISPR Screens. Methods Mol Biol 1712, 203–216 (2018).

50. McAlister, G.C. et al. MultiNotch MS3 enables accurate, sensitive, and multiplexed detection of differential expression across cancer cell line proteomes. Anal Chem 86, 7150–8 (2014).

51. Burgoyne, J.R., Oviosu, O. & Eaton, P. The PEG-switch assay: a fast semi-quantitative method to determine protein reversible cysteine oxidation. J Pharmacol Toxicol Methods 68, 297–301 (2013).

52. Ritchie, T.K. et al. Chapter 11 - Reconstitution of membrane proteins in phospholipid bilayer nanodiscs. Methods Enzymol 464, 211–31 (2009).

53. Punjani, A., Rubinstein, J.L., Fleet, D.J. & Brubaker, M.A. cryoSPARC: algorithms for rapid unsupervised cryo-EM structure determination. Nat Methods 14, 290–296 (2017).

54. Pettersen, E.F. et al. UCSF ChimeraX: Structure visualization for researchers, educators, and developers. Protein Sci 30, 70–82 (2021).

55. Abramson, J. et al. Addendum: Accurate structure prediction of biomolecular interactions with AlphaFold 3. Nature 636, E4 (2024).

56. Cao, H., Li, T., He, J. & Huang, S. BPS2025 - EMReady2: A general model for improving cryo-EM and cryo-ET maps by heterogeneity-aware deep learning. Biophysical Journal 124, 625a (2025).

57. Moriarty, N.W., Grosse-Kunstleve, R.W. & Adams, P.D. electronic Ligand Builder and Optimization Workbench (eLBOW): a tool for ligand coordinate and restraint generation. Acta Crystallogr D Biol Crystallogr 65, 1074–80 (2009).

58. Liebschner, D. et al. Macromolecular structure determination using X-rays, neutrons and electrons: recent developments in Phenix. Acta Crystallogr D Struct Biol 75, 861–877 (2019).

59. Emsley, P., Lohkamp, B., Scott, W.G. & Cowtan, K. Features and development of Coot. Acta Crystallogr D Biol Crystallogr 66, 486–501 (2010).

60. Croll, T.I. ISOLDE: a physically realistic environment for model building into low-resolution electron-density maps. Acta Crystallogr D Struct Biol 74, 519–530 (2018).

61. Williams, C.J. et al. MolProbity: More and better reference data for improved all-atom structure validation. Protein Sci 27, 293–315 (2018).

62. Stierand, K. & Rarey, M. Drawing the PDB: Protein-Ligand Complexes in Two Dimensions. ACS Med Chem Lett 1, 540–5 (2010).

